# The oneirogen hypothesis: modeling the hallucinatory effects of classical psychedelics in terms of replay-dependent plasticity mechanisms

**DOI:** 10.1101/2024.09.27.615483

**Authors:** Colin Bredenberg, Fabrice Normandin, Blake Richards, Guillaume Lajoie

## Abstract

Classical psychedelics induce complex visual hallucinations in humans, generating percepts that are coherent at a low level, but which have surreal, dream-like qualities at a high level. While there are many hypotheses as to how classical psychedelics could induce these effects, there are no concrete mechanistic models that capture the variety of observed effects in humans, while remaining consistent with the known pharmacological effects of classical psychedelics on neural circuits. In this work, we propose the “oneirogen hypothesis”, which posits that the perceptual effects of classical psychedelics are a result of their pharmacological actions inducing neural activity states that truly are more similar to dream-like states. We simulate classical psychedelics’ effects via manipulating neural network models trained on perceptual tasks with the Wake-Sleep algorithm. This established machine learning algorithm leverages two activity phases: a perceptual phase (wake) where sensory inputs are encoded, and a generative phase (dream) where the network internally generates activity consistent with stimulus-evoked responses. We simulate the action of psychedelics by partially shifting the model to the ‘Sleep’ state, which entails a greater influence of top-down connections, in line with the impact of psychedelics on apical dendrites. The effects resulting from this manipulation capture a number of experimentally observed phenomena including the emergence of hallucinations, increases in stimulus-conditioned variability, and large increases in synaptic plasticity. We further provide a number of testable predictions which could be used to validate or invalidate our oneirogen hypothesis.

## 1 Introduction

Classical psychedelics—including psilocybin, mescaline, DMT, and LSD—are a family of hallucinogenic compounds with a common mechanism of action: they are agonists for the 5-HT2a serotonin receptor commonly expressed on the apical dendrites of cortical pyramidal neurons [1] and on parvalbumin (PV) interneurons [2]. These drugs induce numerous effects in human subjects, including: complex visual, auditory, and tactile hallucinations; intense spiritual experiences; long-lasting alterations in mood; changes in personality; and increases in synaptic plasticity [3, 4, 5]. Recently, they have been explored clinically as potential treatments for depression and anxiety [6], as well as PTSD [7].

The 5-HT2a receptor plays a critical role in psychedelic-induced hallucinations. Indeed, behavioral measures of hallucinatory drug effects are induced selectively by cellular membrane-permeable 5-HT2a agonists [8], and perceptual effects of classical psychedelics are largely eliminated by blocking 5-HT2a receptors in the cortex [9, 8] (though 5-HT2a agonists with mixed receptor selectivity are in some cases characterized by primarily non-hallucinatory effects [10, 11]). However, very little is understood about *why* highly structured hallucinations and changes in synaptic plasticity emerge from activating cortical 5-HT2a receptors: to explain this, it is necessary to develop mechanistic theories that are capable of linking changes in neuron-level properties (receptor agonism) to changes in perception and behavior. Psychedelic drug users and therapists have long noted the ‘dream-like’ qualities of psychedelic drug hallucinations, which are realistic but untethered from the external world; this observation leads naturally to speculation that these drugs are ‘oneirogens,’ or dream-manifesting compounds [12]. However, beyond perceptual phenomenology (and some evidence pointing to the effects of psychedelics on sleep cycles [13, 14, 15]), we lack a mechanistic proposal that could explain the similarity between dreams and psychedelic drug experiences. Here, we articulate the ‘oneirogen hypothesis’, which describes one such potential mechanistic explanation. We propose that classical psychedelics induce a dream-like state by shifting the balance between bottom-up pathways transmitting sensory information and top-down pathways ordinarily used to create replay sequences in the brain. Replay sequences have been shown to be important for learning during sleep [16, 17, 18, 19, 20]: we propose that mechanisms supporting replay-dependent learning during sleep are key to explaining the increases in plasticity caused by psychedelic drug administration. In total, our model of the functional effect of psychedelics on pyramidal neurons could provide a explanation for the perceptual psychedelic experience in terms of learning mechanisms for consolidation during sleep [21], and cortical ‘replay’ phenomena [22, 23, 24, 25, 26, 27, 28, 29, 30, 31, 32].

To explore the oneirogen hypothesis concretely, we use the aptly named Wake-Sleep algorithm [33], which has historically been used to train artificial neural networks (ANNs) that possess both a bottomup “recognition” pathway and a top-down “generative” pathway to learn a representation of incoming sensory data. It enables unsupervised learning in ANNs by alternating between periods of “waking perception” (wherein bottom-up recognition pathways drive activity) and “dreaming sequences” (wherein top-down generative pathways drive activity). With these alternate periods of distinct activity, connectivity parameters in each pathway are adjusted to match the activity of the opposite pathway. This way, the top-down pathway learns to generate activity consistent with that induced by sensory inputs, and the bottom-up pathway learns better representations thanks to generated activity.

In this work, we show that within a neural network trained via Wake-Sleep, it is possible to model the action of classical psychedelics (i.e. 5-HT2a receptor agonism) by shifting the balance during the wake state from the bottom-up pathways to the top-down pathways, thereby making the ‘wake’ network states more ‘dream-like’. Specifically, we model the effects of classical psychedelics by manipulating the relative influence of top-down and bottom-up connections in neural networks trained with the Wake-Sleep algorithm on images. Doing so, we capture a number of effects observed in experiments on individuals under the influence of psychedelics, including: the emergence of closed-eye hallucinations, increases in stimulus-conditioned variability, and large increases in synaptic plasticity. This data suggests that the oneirogen hypothesis may indeed help to explain why 5-HT2a agonists have the functional effects that they do. We subsequently identify several testable predictions that could be used to further validate the oneirogen hypothesis.

## 2 Results

### Mapping the Wake-Sleep algorithm onto cortical architecture

The Wake-Sleep algorithm allows ANNs to optimize a global, unsupervised objective function for sensory representation learning— the Evidence Lower Bound (ELBO)—through local synaptic modifications to a bottom-up recognition pathway and a top-down generative pathway. As a precursor to the variational autoencoder [34, 35], the Wake-Sleep algorithm provides a mechanism for learning a probabilistic latent representation **r** responding to incoming sensory stimuli **s**, which obeys representational characteristics that are ideal for a neural system (e.g. sparsity and metabolic efficiency [36], compression and coding efficiency [37, 38], or disentanglement [39, 40]). To do this, Wake-Sleep optimizes the ELBO through an approximation of the Expectation Maximization (EM) algorithm [41] to train the two pathways (Figure 1a).^1^

**Figure 1.**
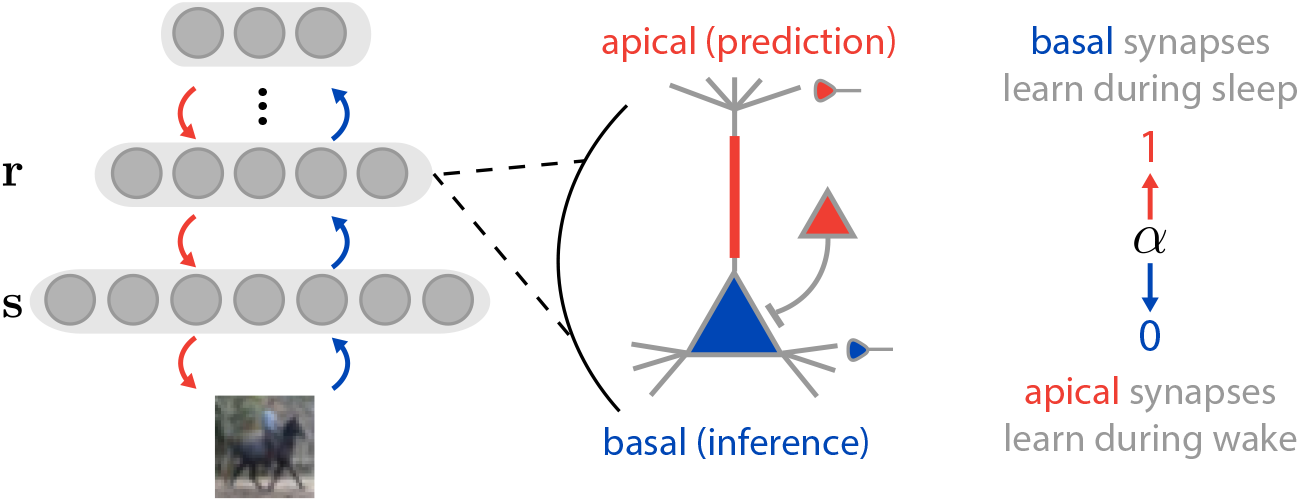
Mapping the Wake-Sleep algorithm onto cortical architecture. Left: Network architecture. We model early sensory processing in the cortex with a multilayer network, **r**, receiving stimuli **s**. Center: individual pyramidal neurons receive top-down inputs (red) at the apical dendritic compartment, and bottom-up inputs at the basal dendritic compartment (blue). 5-HT2a receptors are expressed on the apical dendritic shaft (red bar), and on PV interneurons (red triangle); both sites may play a role in gating basal input. Right: Over the course of Wake-Sleep training, basal inputs dominate activity during the Wake phase (*α* = 0) and are used to train apical synapses, whereas apical inputs dominate activity during the Sleep phase (*α* = 1) and are used to train basal synapses.

Notably, the Wake-Sleep algorithm requires two phases of activity (i.e. “Wake” and “Sleep”), where the network phase is controlled by a global state variable *α ∈*[0, 1] that regulates the balance between the bottom-up and top-down pathways. In the Wake phase (*α* = 0), the network processes real sensory stimuli drawn from the environment, and network activity is sampled based on the bottom-up inputs (corresponding to the approximate inference distribution). In the Sleep phase (*α* = 1), the network internally samples neural activity from its generative model, which then produces generated activity in the stimulus layer **s**. We use this structure of the Wake-Sleep algorithm as a concrete model to express oneirogen hypothesis. Specifically, we use changes to the value of *α* as a means of modeling a 5-HT2a agonist-induced shift to a more dream-like state, as we detail below.

Within the Wake-Sleep algorithm, neurons alternate between ‘Wake’ and ‘Sleep’ modes, where activity during each mode is dominated by the bottom-up and top-down pathways, respectively. We can determine the neural activity for a given intermediate layer *l* with the following equation:

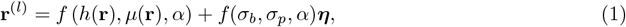

where *h*(**r**) defines bottom-up input, *µ*(**r**) defines top-down input, *f* (*h, µ, α*) is any interpolation function such that *f* (*h, µ*, 0) = *h* and *f* (*h, µ*, 1) = *µ, σ*_*b*_ and *σ*_*p*_ define the bottom-up and top-down activity standard deviations, and ***η*** *∼𝒩* (0, 1) adds random noise to the neural activity (see Methods for more detail). Here, for notational conciseness we treat **r** as a concatenated vector of all **r**^(*l*)^ vectors from each layer. This equation means that *α* controls whether bottom-up inputs or top-down inputs control the dynamics of individual neural units.

Thus, as *α* moves from a value of 0 to a value of 1, the activity of the neurons shifts from being driven by the bottom-up recognition pathway to being driven by the top-down generative pathway. How could this occur in the brain? Realistically, each neuron in cortex would have its own *α* variable defining the relative influence of top-down and bottom-up inputs on its spiking activity; here, for simplicity, we will assign the entire network a single *α* value reflecting the ‘mean’ relative top-down/bottom-up influence averaged across neurons, as determined by the network state (Wake, Sleep, dose-dependent psychedelic administration). In the cortex, excitatory pyramidal neurons receive inputs from distinct sources: inputs that are from ‘higher order’ cortical areas target the apical dendrites, whereas inputs that are from ‘lower order’ cortical or sensory subcortical areas target the basal dendrites [43]. Thus, we can capture the core idea behind the oneirogen hypothesis using the Wake-Sleep algorithm, by postulating that the bottom-up basal synapses are predominantly driving neural activity during the Wake phase (when *α* is low), while top-down apical synapses are predominantly driving neural activity during the Sleep phase (when *α* is high; Figure 1) [44]; this is in agreement with several recent theoretical studies that have proposed that apical dendrites could serve as a site for integrating top-down learning signals [45, 46, 47, 48, 49, 50], particularly those which propose that the top-down signal corresponds to a predictive or generative model of neural activity [51, 52]. This proposed change in *α* does indeed appear to occur during both slow-wave (SW) [53, 54] and rapid eye movement (REM) [55, 56, 57] sleep, where apical dendritic inputs have been observed to exert increased influence on neural activity that is critical for plasticity induction and consolidation of learned behaviors; during REM sleep, this increased influence has been shown to be mediated by potentiation of basal dendrite-targeting PV inhibitory interneurons [57].

Next we ask: can we model the effects of classical psychedelics in terms of changes in *α*? Notably, 5-HT2a receptors are expressed in the apical dendrites of pyramidal neurons [1] and PV interneurons [2], and have an excitatory effect that positively modulates glutamatergic transmission due to apical dendritic inputs [58, 59]; further, classical psychedelic administration has been shown to have an inhibitory effect on glutamatergic transmission due to basal dendritic inputs [60]. These data suggest that 5-HT2a agonists could have a push-pull effect on cortical pyramidal neurons, increasing the relative influence of apical dendrites and decreasing the relative influence of basal dendrites [61] in much the same way as has been observed during SW and REM sleep. Hence, we can model these effects by increasing the *α* value in a Wake-Sleep trained network, and then ask whether the networks exhibit other phenomena that match the known impact of classical psychedelics on neural activity. We note that with this mapping of the Wake-Sleep algorithm to models of basal and apical processing, synaptic modifications at both apical and basal synapses correspond to minimizing a local prediction-error between top-down and bottom-up inputs (see Methods).

### Modeling Hallucinations

To see whether a transition from waking to a more dream-like state would induce hallucinatory effects in our model, we trained multilayer neural networks with branched dendritic arbors (see Methods) on the MNIST digits dataset [62] using the Wake-Sleep algorithm and subsequently simulated hallucinatory activity by varying *α* (see Methods; Eq. (8)). We could visualize the effects of our simulated psychedelic with snapshots of the stimulus layer **s** at a fixed point in time for various values of *α* (Figure 2; see Supplemental Materials for videos). As *α* increased, we observed that network activity gradually deformed away from the ground-truth stimulus in a highly structured way, adding strokes to the original digit that were not originally present. At the highest values of *α* tested, we found that network states were wholly divorced from the ground-truth stimulus, but retained many characteristics of the MNIST digits on which the network was trained (e.g. smooth strokes and the rough form of digits). These results emphasize that hallucinations induced by a shift to a more dream-like state in these models are heavily influenced by the training dataset, which for an animal would correspond to the statistics of the sensory environment in which it learns its sensory representation. To emphasize this point, we further trained our networks on the CIFAR10 natural images dataset [63] (Figure 2c), to provide an example of a more naturalistic training dataset. In this case, our model was not powerful enough to reproduce realistic natural images—instead, we found that our modeled hallucinatory activity corresponded to ‘ripple’ effects, which are similar to the ‘breathing’ and ‘rippling’ phenomena reported by psychedelic drug users at low doses [3].

**Figure 2.**
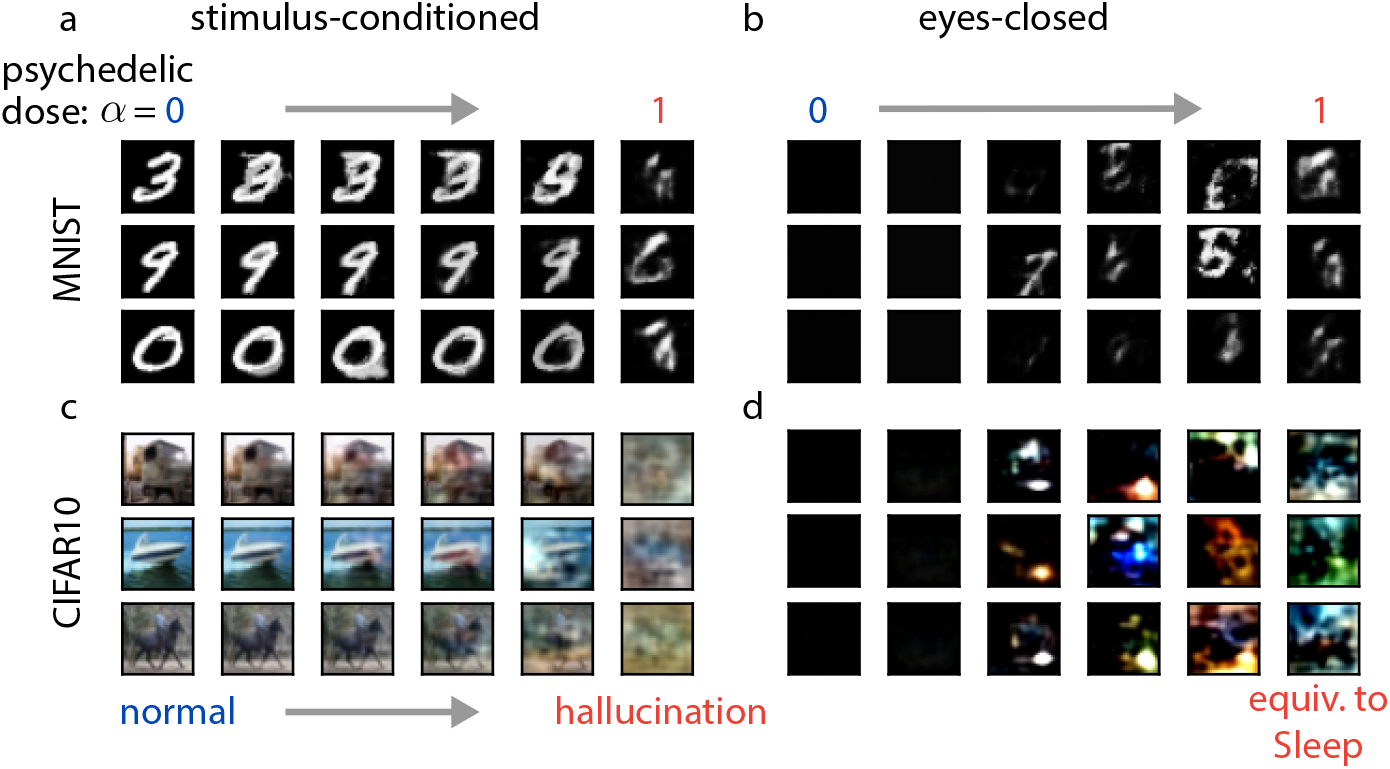
Visualizing the effects of psychedelics in the model. We model the effects of classical psychedelics by progressively increasing *α* from 0 to 1 in our model, where *α* = 1 is equivalent to the Sleep phase. We visualize the effects of psychedelics on the network representation by inspecting the stimulus layer **s. a)** Example stimulus-layer activity (rows) in response to an MNIST digit presentation as psychedelic dose increases (columns, left to right). **b)** Same as (a) but for ‘eyes-closed’ conditions where an entirely black image is presented. **c-d)** Same as (a-b), but for the CIFAR10 dataset.

These simulations were produced with a complex, multicompartmental neuron model; however, we found similar results with two alternative network architectures, one with within-layer recurrence (Supplemental Figure S1a) and one which used a simpler single compartment neuron model (Supplemental Figure S1b). We found that our single compartment model produced qualitatively less realistic generated images than the multicompartment and recurrent models, justifying our use of the more complex models (Supplemental Figure S2). To demonstrate the importance of a learned top-down pathway to produce complex, structured hallucinations in the earliest layers of our network, we generated model hallucinations from two control networks: an untrained model and a trained network where psychedelic activity was alternatively modeled by a simple increase in the variance of individual neurons (we will refer to this latter control as the noise-based hallucination protocol). We found that hallucinations under these control conditions resembled additive white noise, rather than structured digit-like shapes (Supplemental Figure S1c-d).

Psychedelic drug users also report observing the emergence of hallucinations while their eyes are closed [3]. Interestingly, we found that our model recapitulated these phenomena: as *α* increased, networks trained on MNIST gradually began revealing increasingly complex and digit-like patterns (Figure 2b), whereas CIFAR10-trained networks again predominantly produced ‘ripple’ hallucinations (Figure 2d).

### Effects of psychedelics on single neurons

Having recapitulated hallucinatory phenomena in stimulus space, we next explored how our proposed mechanism affected neural activity in our network model, in order to establish markers that could be used to experimentally validate or invalidate the oneirogen hypothesis. To start, we investigated the effects of learning and psychedelic drug administration on the activity of single neurons in the model. As noted previously, the learning algorithm used here trains synapses so that top-down inputs to apical dendritic compartments match bottom-up inputs to basal dendritic compartments. As a consequence, we observed that after training, inputs to apical and basal dendritic compartments were much more correlated on the same neuron than they were for random neurons (Figure 3a), which was not observed in untrained models (Supplemental Figure S3a). This form of strongly correlated tuning has been observed in both cortex and the hippocampus [64, 65].

**Figure 3.**
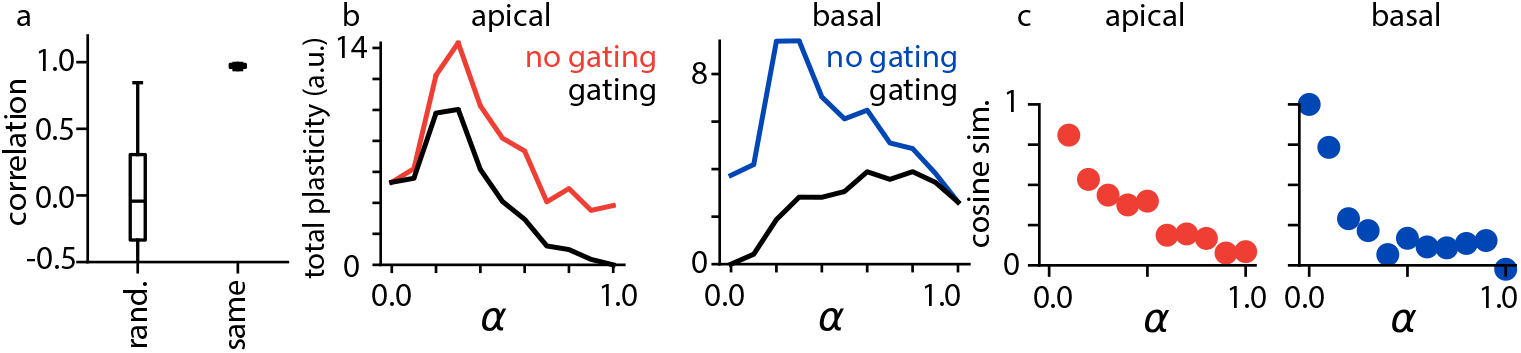
Effects of psychedelics on single model neurons. **a)** Correlations between the apical and basal dendritic compartments of either the same network neuron or between randomly selected neurons. **b)** Total plasticity for apical (left) and basal (right) synapses as *α* increases in the model when plasticity is either gated or not gated by *α*. Error bars indicate +/-1 s.e.m. **c)** Cosine similarity between plasticity induced under psychedelic conditions compared to baseline for apical (left) and basal (right) synapses.

There are many indicators that psychedelic drug administration in humans and animals can induce marked, long-lasting changes in behavior, as well as large increases in synaptic plasticity [4, 66, 67, 8, 5]. In Wake-Sleep learning, apical synapses learn during the Wake phase, whereas basal synapses learn during the Sleep phase—thus, plasticity at apical synapses is gated by (1*− α*), whereas plasticity at basal synapses is gated by *α* (see Methods). However, learning is still theoretically possible without this explicit gating, though it may be noisier and less efficient; furthermore, it is conceivable that classical psychedelics could increase the relative influence of apical inputs on the activity of a neuron without affecting this gating mechanism. As a consequence, we modeled the dose-dependent effects of psychedelics on plasticity both with and without gating (Figure 3b). Consistent with recent experimental results, for intermediate doses we found large increases in plasticity at both apical and basal synapses under *both* conditions, where plasticity was measured as a mean change in normalized synaptic strength across weight parameters in our network (see Methods). In our model, we found that the total evoked plasticity peaked at roughly *α* = 0.5; we further found that if gating was affected by psychedelics, apical plasticity would eventually be quenched at very high drug doses. We also found that plasticity induced by psychedelic drug administration gradually became unaligned from the weight updates that would have occurred in the absence of the drug (Figure 3c), indicating that these results were not simply due to modulation of the effective learning rate of the underlying plasticity. Rather, as has been suggested by other theoretical studies [68], plasticity in the model likely increased because aberrant hallucinatory activity pulled the learning mechanism out of a local optimum in which plasticity was minimal, producing much more plasticity across the network. Importantly, we observed these increases in plasticity in all network architectures and training datasets we explored, including for our noise-based hallucination protocol (Supplemental S4), demonstrating that changes in apical dendritic influence within a Wake-Sleep learning framework are sufficient, but *not necessary* to induce increases in synaptic plasticity: for trained networks, it would seem that even simple increases in neural variability can have similar effects.

### Effects of psychedelics on neural variability

Having observed that increasing our modeled drug dosage caused heightened fluctuations and deviations from the ground-truth stimulus in the sensory layer of our network (Figure 2), we next investigated whether variability was affected at the level of individual neurons in higher layers of the model. Indeed, we found that for a fixed stimulus, neural variability increased markedly as the simulated psychedelic drug dose increased (Figure 4a). This result is consistent with the data supporting the Entropic Brain Theory [69, 70, 71, 72], in which neural activity in resting state fMRI recordings becomes increasingly ‘entropic’ (i.e. variable) under the influence of psychedelics; however, it is important to note that our noise-based hallucination protocol also produced these effects (Supplemental Figure S5a). Though most experimental data supporting the Entropic Brain Theory is taken from recordings with relatively poor spatial resolution, averaging activity over large cortical areas, our model predicts that this increase in variability should be reflected at the level of individual neurons; this increase in variability after psychedelic administration has been recently observed in auditory cortical neurons for active mice [73], but whether this phenomenon is general across tasks and cortical areas remains to be seen. We further found that this increase in variability corresponded to a decrease in ability to identify the stimulus being presented to the network: we trained a classifier to identify which MNIST digit was presented to our networks on Wake neural activity (see Methods), and found that the accuracy of our classifier decreased (Figure 4b) while the output variability of the classifier increased (Figure 4c) in response to drug administration.

**Figure 4.**
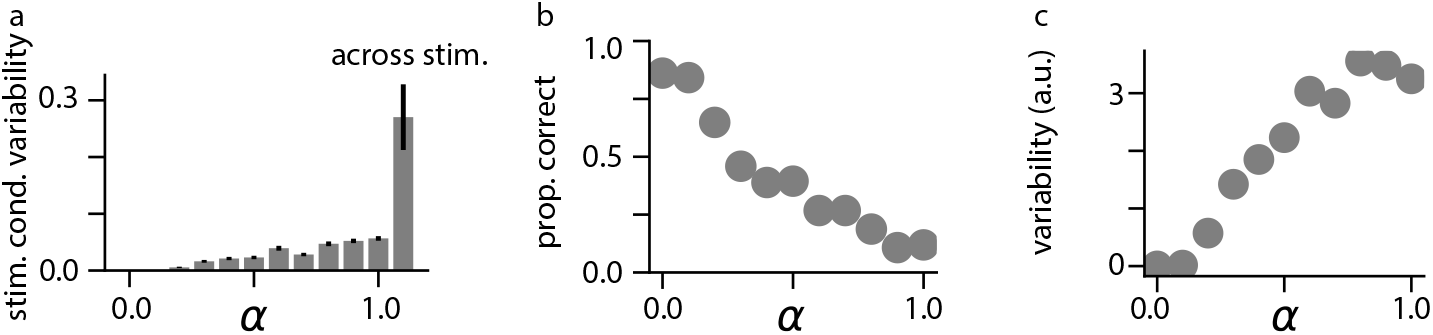
Effects of psychedelics on neural variability. **a)** Stimulus-conditioned variability for neurons in the network as *α* increases, as compared to variability in neural activity across stimuli (rightmost bar). Error bars indicate +/-1 s.e.m. **b)** Proportion correct for a classifier trained to detect the label of presented MNIST digits as *α* increases. **c)** Variability in the logit outputs of the trained classifier as *α* increases.

Within our model, this increase in variability is quite sensible: in the ordinary Wake state, neural activity is constrained to correspond to the singular sensory stimulus being presented, whereas during Sleep states, neural activity is completely unconstrained by any particular sensory stimulus, reflecting instead the full distribution of *possible* sensory stimuli. As increasing *α* in our model interpolates between Wake and Sleep states, we can expect intermediate values of *α* to produce network states which are less constrained by the particular sensory stimulus being presented, reflected in increased neural variability.

### Network-level effects of psychedelics

We next investigated the effects of psychedelics on networklevel and inter-areal dynamics within our model. We first identified an important negative result: the pairwise correlation structure between neurons was largely preserved across psychedelic doses (Figure 5a-b), as was the effective dimensionality of population activity (Figure 5c). This was sensible, because a network that has been well-trained with the Wake-Sleep algorithm will have the same marginal distribution of network states in the Wake mode as in the Sleep mode—thus, pairwise correlations between neurons should also not differ (as measures of the second order moments of the marginal distribution). We found empirically that even for intermediate values of *α* in which activity is a mixture of Wake and Sleep modes, these correlations are largely unchanged; in contrast, we observed large changes in correlation structure for untrained networks, and increases in effective dimensionality for both untrained networks and for our simple noise-based hallucination protocol, suggesting that these results are more specific to our trained models in which hallucinations are caused by an increase in apical dendritic influence (Supplemental Figure S6a-b). Interestingly, these results are consistent with a recent study that has shown only minimal functional connectivity and effective dimensionality changes in task-engaged humans being presented audiovisual stimuli under the influence of psilocybin [72].

**Figure 5.**
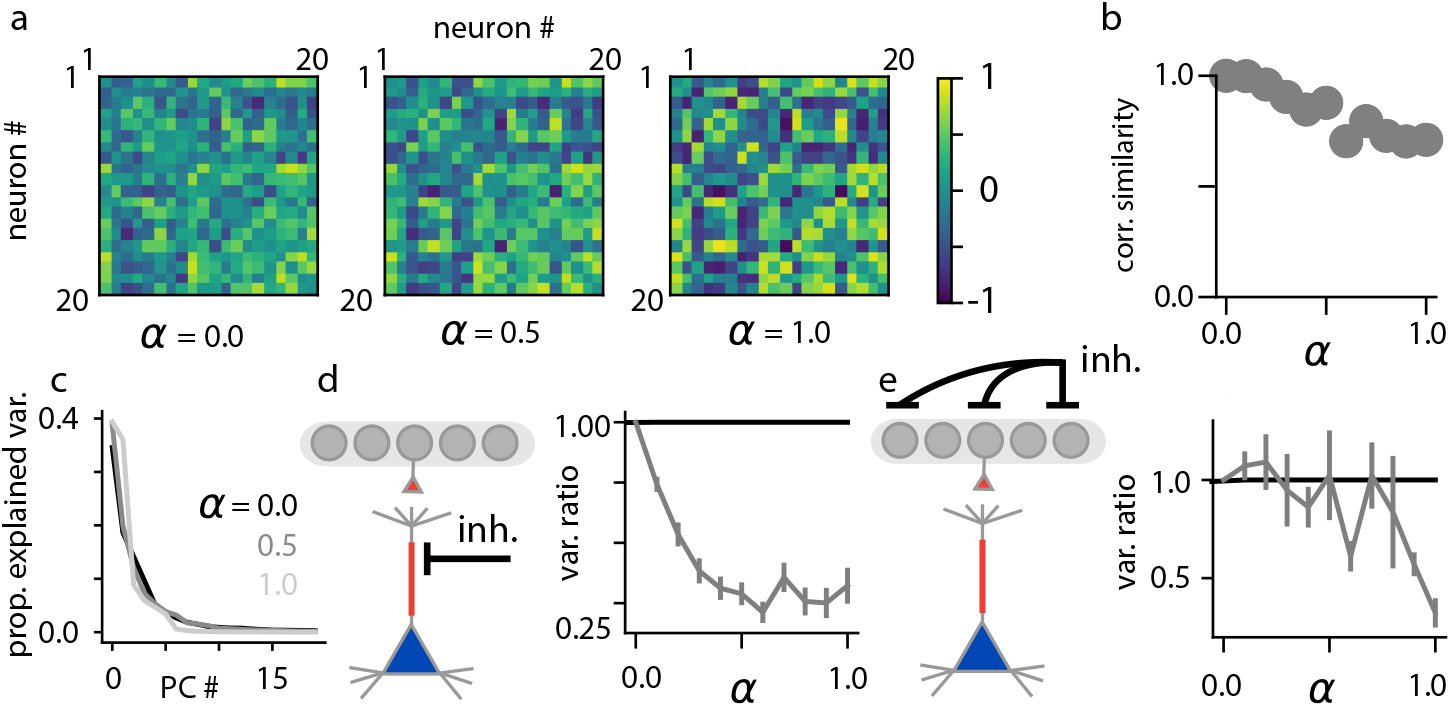
Network-level effects of psychedelics. **a)** Pairwise correlation matrices computed for neurons in layer 2 across stimuli for *α* = 0 (left), *α* = 0.5 (center), and *α* = 1.0 (right). **b)** Correlation similarity metric between the pairwise correlation matrices of the network in the absence of hallucination (*α* = 0) as compared to hallucinating network states (*α >* 0). **c)** Proportion of explained variability as a function of principal component (PC) number for *α ∈ {* 0, 0.5, 1*}*. **d)** Ratio of across-stimulus variance in individual stimulus layer neurons when the apical dendrites have been inactivated, versus baseline conditions across different *α* values. **e)** Ratio of across-stimulus variance in individual neurons in the stimulus layer when neurons at the deepest network layer have been inactivated, versus baseline conditions across different *α* values. Error bars indicate +/-1 s.e.m.

However, though the pairwise correlations between single neurons are largely preserved, the causal influence between lower and higher layers of our model network changes considerably both during hallucination and Sleep modes. Because psychedelic drug administration increases influence of apical dendritic inputs on neural activity in our model, we found that silencing apical dendritic activity reduced across-stimulus neural variability more as the psychedelic drug dose increases (Figure 5d). Further, we found that as *α* increased, inactivating the deepest network layer induced a large reduction in variability in the stimulus layer relative to baseline (Figure 5e), revealing that within our model, increases in top-down influence are responsible for much of the observed stimulus-conditioned variability at larger drug doses. These inactivations had no impact on neural variability in our noise-based hallucination protocol, but were observed for all network architectures and datasets that we tested in which hallucinations were caused by an increase in apical dendritic influence (Supplemental Figure S6), suggesting that these results are quite specific to our model. Further, these inactivations have not yet been performed in animals, and consequently constitute a critical testable prediction of our model.

### Modeling hallucinations in large-scale pretrained networks

While our trained model is capable of capturing several effects of classical psychedelics, it also has a clear limitation: our top-down generative model does not have sufficient expressive power to induce complex hallucinations of naturalistic stimuli, producing instead ‘ripples,’ or ‘breathing’ effects that preserve lower-order statistical features of the input data (Figure 5b). While psychedelic drug users do report these phenomena, they also report observing much more complex hallucinations, including of people, animals, and scenes [74, 75].

Generative models trained through backpropagation have been much more successful in producing more complex generated sensory stimuli [35, 34, 76], and furthermore, hierarchical variational autoencoder models have a nearly identical top-down/bottom-up model architecture as our Wake-Sleep-trained networks [77, 78]. Therefore, to see whether our proposed mechanism would induce complex, structured hallucinations in more powerful models, we induced hallucinations in Very Deep Variational Autoencoder (VDVAE) models [79] that were pretrained through backpropagation on a large natural images dataset, Tiny ImageNet [80], and a large corpus of human faces, FFHQ-256 [81]. These models have a few key differences compared to our Wake-Sleep-trained models: 1) they are trained through backpropagation, which is well-known to be biologically implausible [82]; 2) they exploit parameter sharing across spatial positions in convolutional layers for increased data efficiency during training, at the expense of further biological realism [83]; 3) the ‘Wake’ stage inference process of these models incorporates inputs from both bottom-up *and* top-down sources, which both improves performance [77] and is more biologically realistic [84, 43]; 4) the models are trained on more complex, higher-resolution datasets. Finally, to induce more ‘abstract’ hallucinations, we increased the *α* parameter in these models selectively for higher layers of the network, whereas for the Wake-Sleep-trained models we increased *α* evenly across layers (see Methods). Combined, these differences make for an effective model of high-level hallucination effects, at the expense of some biological realism.

We found that hallucinations generated by these pretrained models were much richer and more complex: increasing *α* in the Tiny ImageNet VDVAE caused the emergence of textural patterns and geometric shapes, while the FFHQ-256 VDVAE caused increasingly bizarre changes in facial features (Fig. 6). Both models were also capable of reproducing closed-eyes hallucinations (Fig. S7), where the content of these hallucinations was shaped by their respective training datasets.

**Figure 6.**
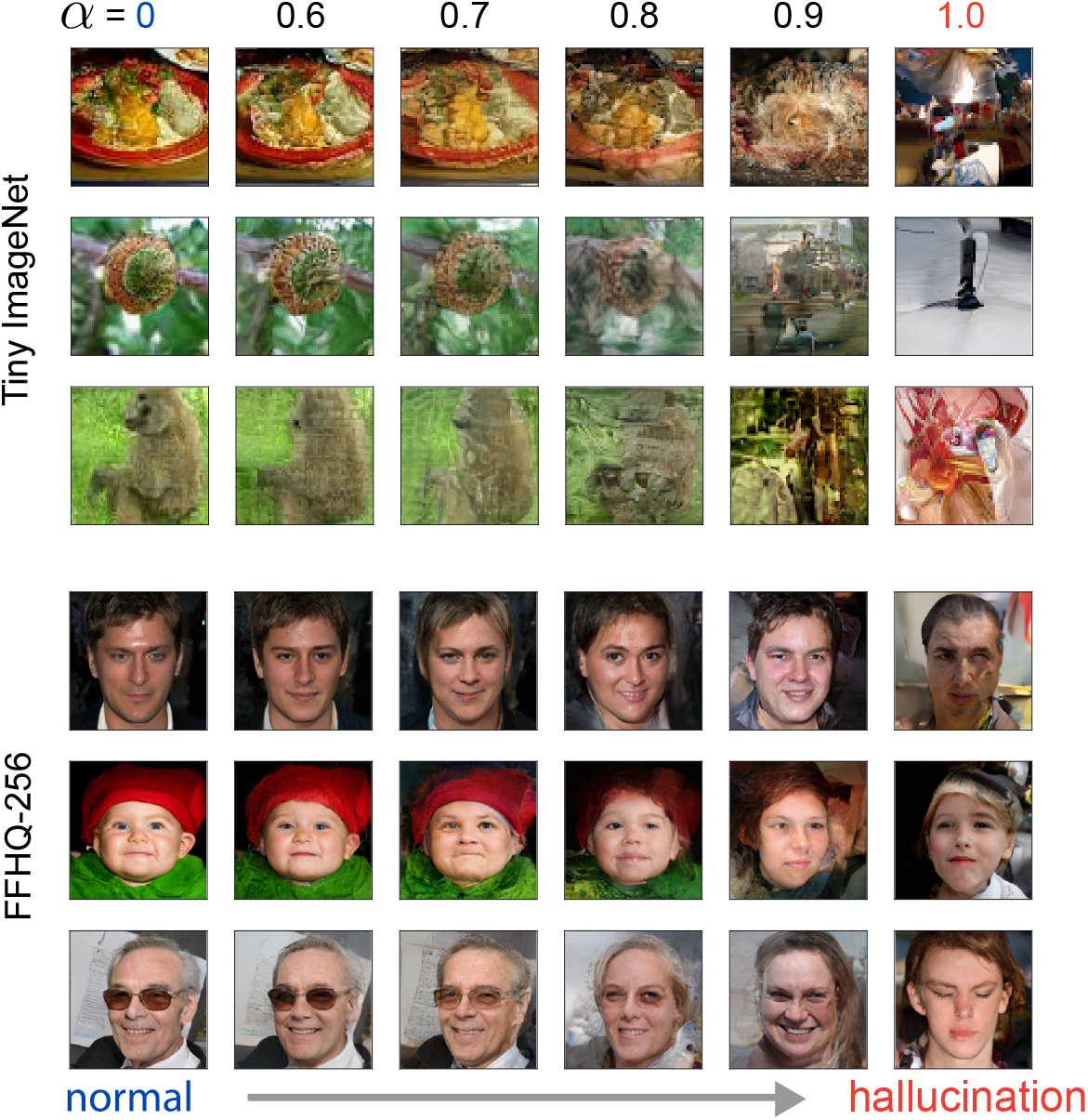
Visualizing the effects of psychedelics in pretrained VDVAE models. Decoded outputs of a pretrained VDVAE model trained on Tiny ImageNet (Top) and FFHQ-256 (Bottom) based on hallucinations generated in the top 35 layers of the model. Image samples vary along rows, and hallucination intensity, parameterized by *α*, increases along columns.

To investigate the nature of hallucinations generated by the Tiny ImageNet VDVAE, we examined the Laplacian pyramid of decoded hallucination images at varying *α* values (Fig. S8a). Essentially, a Laplacian pyramid decomposes an image into levels of decreasing resolution features, with each level encoding the residual produced by downsampling to the next-lowest resolution (level 0 corresponds to the base 64x64 pixel image, while level 5 corresponds to a 4x4 reduced-resolution set of features). We found that low-level pyramid features varied considerably at low *α* levels, while high-level pyramid features did not begin to vary until higher *α* doses (Fig. S8b). This suggests that hallucinations within our model obey a fine-to-coarse structure, where low-dose hallucinations are confined to high-frequency, spatially localized changes, and progressively increasing doses begin to cause variations in more global image features.

Lastly, we were able to replicate our previous network-level results on the Tiny ImageNet VDVAE. We found that increasing psychedelic dose *α* caused an increase in stimulus-conditioned variance within the model (Fig. S8c), and that across-stimulus correlation structure between network units was largely preserved across doses (Fig. S8d). Further, we found that the ratio of before- and after-inactivation across-stimulus variance decreased as the psychedelic dose *α* increased (though somewhat paradoxically, inactivation caused an *increase* in variance for *α* = 0, likely due to the influence of top-down inputs during inference for this model). Combined, these results show that key testable predictions from our Wake-Sleep-trained model are preserved in the VDVAE, while this latter model is capable of producing some of the more complex hallucinations characteristic of psychedelic experience.

## 3 Discussion

### Experimental results captured by our model

In this study, we have examined a hypothetical mechanism explaining how the 5-HT2a receptor agonism of classical psychedelics could induce the highly structured hallucinations reported by people who have consumed these drugs. Specifically, we have explored the ‘oneirogen hypothesis’, which postulates that 5-HT2a agonists have the effects that they do because they shift the neocortex to a more dream-like state, wherein activity is more strongly driven by top-down inputs to apical dendrites than normally occurs during waking. To provide a concrete model to explore the ‘oneirogen hypothesis’ we used the classic Wake-Sleep algorithm, which learns by toggling between a Wake phase, where activity is driven by bottom-up sensory inputs, and a Sleep phase, where activity is driven by top-down generative signals. We modeled the ‘oneirogen hypothesis’ by simulating psychedelic administration as an increase in a neuronal state variable (*α*) that switches neural activity between these two phases, such that the simulated psychedelic caused the network to enter a state somewhere between the Wake and Sleep phases, making activity during the Wake phase less tied to actual sensory inputs by increasing the relative influence of the top-down, apical compartment in the models (depending on the “dosage”). This formulation is consistent with anatomical wiring data [43], as well as several recent theoretical studies which propose a specialized learning role for top-down projections to the apical dendrites of pyramidal neurons [45, 46, 47, 48, 49, 50]. It is also consistent with the known cellular mechanism of action of classical psychedelics [1, 58, 59, 9] and experiments that demonstrate a reduced responsivity to bottom-up stimuli in cortex after psychedelic drug administration [85, 86, 87]. Using this model, we were able to produce both stimulus-conditioned and “closed-eye” hallucinations that are consistent with the low-level effects reported by psychedelic drug users [3], and we were also able to recapitulate the large increases in plasticity observed at both apical and basal synapses at moderate psychedelic doses [4].

Our model uses a particular functional form of synaptic plasticity at both apical and basal synapses, reminiscent of the classical delta rule [88], which seeks to minimize a prediction error between inputs in apical and basal synapses. There are many theoretical models of learning that propose similar forms of plasticity [46, 47, 51], so while this plasticity is a necessary prediction of our model, it is not *sufficient* to validate it. Experimentally, plasticity dynamics which could, theoretically, minimize such a prediction error have been observed in cortex [89, 90]; we found that plasticity rules of this kind induce strong correlations between inputs to the apical and basal dendritic compartments of pyramidal neurons, which has been observed in both the hippocampus and cortex [64, 65]. Psychedelic administration within our model induced large increases in plasticity, which has also been observed experimentally [4, 5]. Within our model, this plasticity should not be interpreted as ‘learning,’ since it arises from aberrant network activity and does not necessarily produce behavioral or perceptual improvements; it is likely closer to ‘noise,’ that may still be useful for helping neural networks escape from local minima in the loss optimization landscape for synaptic weights, with possible implications for individuals suffering from post-traumatic stress disorder, early life trauma, or the negative effects of sensory deprivation. Further work will be required to analyze the relationship within our model between psychedelic dosage, usage frequency, and the long-term stability of learned representations in neural networks.

Interestingly, we also found that increasing the influence of apical dendrites in the model increased stimulus-conditioned variability in our individual neurons. In cortex, this effect has recently been shown at the level of single auditory neurons [73]; further, there have been numerous studies reporting similar increases in asynchronous variability [71] (or, analogously, sample entropy [70] and Lempel-Ziv complexity [91]) in resting-state human brain recordings, previously modeled using Entropic Brain Theory. This theory proposes that many of the effects of classical psychedelics on perception and learning can be explained in terms of increases in variability induced by drug administration (e.g. the increase in variability could introduce novel patterns of thinking, or perturb learning to allow it to break out of ‘local minima’). Our results are broadly consistent with this perspective, to which we have added explanatory layers that are both normative and mechanistic [92, 93]: namely, we speculate that this variability under ordinary conditions results from an ethologically important mechanism underlying generative replay for unsupervised learning during sleep or quiescence, and we propose that mechanistically this increase in variability is caused by the increased influence of top-down synapses that are not tied to incoming sensory stimuli. Alternatively, such entropy increases could be caused by increases in attention or self-reflective thought, as supported by recent studies showing that task engagement significantly attenuates psychedelic-induced entropy increases [72]; though our model does not include cognitive or attention components, such an interpretation is potentially consistent with and complementary to our framework.

### Testable predictions

While our results are broadly consistent with existing experimental evidence, there are many unconfirmed aspects of our model which could be tested to validate or invalidate it (summarized in Table 1). As mentioned in the previous section, our model predicts that *single neurons* should increase variability in response to psychedelic drug administration in any cortical area affected by psychedelic drugs, an effect that has not yet been investigated systematically throughout cortex or across task conditions. Second, we propose that psychedelic drugs should not push network dynamics into wildly different operating regimes than normal wakefulness, beyond any differences observed between wakefulness and replay (dreams) during sleep. In particular, we found that our simulated psychedelic drug administration did not perturb pairwise correlations between neurons within local circuits when averaged across an ecologically representative set of stimuli.

**Table 1:** Summarizing testable predictions of the ‘oneirogen hypothesis’. Models: OH - oneirogen hypothesis; EC - Ermentrout & Cowan [94]; REBUS - Relaxed Beliefs Under Psychedelics [69]; DD - DeepDream [95]. Key: ✓- model is consistent with the prediction; ✓-model is inconsistent with the prediction; n/a - model is neither inconsistent nor consistent with the prediction.

| Testable Predictions | OH | EC | REBUS | DD |
| --- | --- | --- | --- | --- |
| 1. Psychedelic administration increases stimulus-conditioned variability of neurons. | ✓ | ✓ | ✓ | ✓ |
| 2. Psychedelic administration preserves pairwise across-stimulus correlations between neurons. | ✓ | ✗ | ✗ | ✗ |
| 3. Silencing apical dendritic compartments decreases neural variability more after psychedelic administration. | ✓ | n/a | n/a | n/a |
| 4. Silencing higher-order cortical areas affects lower-order cortical activity more after psychedelic administration. | ✓ | ✗ | ✗ | ✓ |
| 5. Psychedelic drug effects are mediated by the same circuitry responsible for inducing generative replay dynamics in cortex. | ✓ | ✗ | ✗ | ✗ |

Within our model, psychedelic drug administration is expected to increase the relative influence of top-down projections. This prediction appears to be supported by slice experiments [58, 59, 60], but to our knowledge this change in functional connectivity has not yet been shown via *in vivo* manipulations. This could be explored experimentally in several ways: first, we have shown that apical dendrite-targeted silencing experiments can identify the amount of influence apical dendritic inputs exert on neuronal dynamics; second, we have shown that increases in top-down influence can in principle be identified with interareal silencing experiments. We caution that interpreting results in this second vein may be difficult, as establishing a clean distinction between a ‘higher order’ and ‘lower order’ cortical area may be much more difficult in a densely recurrent system such as the brain, compared to our simplified and fully observable network model.

Interestingly, if psychedelic drugs are genuinely co-opting circuitry ordinarily reserved for generative replay during periods of offline quiescence or sleep, we would expect that the same changes in functional connectivity observed during psychedelic drug administration would also occur during periods of replay. Replay been observed and dreams have been documented during both SW [23, 25] and REM [31, 32] sleep, with REM dreams exhibiting greater degrees of bizarreness, possibly indicating a more ‘generative’ form of replay [96]. During SW sleep, increased top-down influence has been observed from secondary motor cortex to primary somatosensory cortex [54], and from hippocampus to prefrontal cortex [25]; however, it should be noted that increased hippocampal-to-prefrontal functional coupling was not observed after classical psychedelic administration [97]. During REM sleep, increased top-down influence (or apical dendritic influence) has been observed in prefrontal, visual [56], and motor [55] cortices, with some top-down inputs originating from higher-order thalamic nuclei [57, 98]; similarly, multiple non-invasive imaging studies have observed increases in top-down functional coupling from higher-order thalamic nuclei after psychedelic administration [99, 100]. Therefore, increases in top-down coupling appear broadly consistent between REM sleep and classical psychedelic administration, while psychedelic states appear inconsistent with the hippocampal-cortical coupling during SW sleep; this latter result could potentially be explained in terms of a recent complimentary learning systems model [101], in which SW sleep is responsible for orchestrating hippocampus-cortex-coupled *episodic* replay while REM sleep is responsible for orchestrating hippocampus-cortex-decoupled *generative* replay, but more experiments and theoretical work will likely be necessary to fully characterize this additional complexity. Given these data, it seems as though REM sleep replay is a moderately stronger candidate for sharing a mechanism of action with classical psychedelics, though it remains possible that replay events during SW sleep occur via a similarly shared mechanism.

To summarize, though we have provided a candidate explanation for several of the hallucinatory effects of psychedelic drugs with a model that displays a strong correspondence with existing empirical evidence, our model rests on a number of testable assumptions. Our goal here has been to articulate these assumptions as clearly as possible, to facilitate experimental efforts to test them.

### Comparisons to alternative models

Here, we review prominent existing hypotheses as to how psychedelic drugs could induce hallucinations in neural networks and compare to our model (summarized in Table 1). The first alternative proposed that incredibly complex, geometric patterns formed by DMT administration could be attributed to pattern-formation effects in visual cortex caused by a disruption of the balance between excitation and inhibition in locally-coupled topographic recurrent neural networks [94, 102]. Our work differs from this approach in several respects. First, rather than disrupting E-I balance, we propose that psychedelics increase the relative influence of apical dendrites and top-down projections on the dynamics of neural activity. Second, though their model is able to generate geometric patterns, it is not able to generate patterns that are statistically related to the features of the sensory environment (e.g. MNIST digits). Lastly, for simplicity we avoided including topographic (or convolutional) recurrent connectivity in our model; however, it would be a very fruitful direction for future research to extend our work to generative modeling of temporal video sequences, as in [103, 104]. With such a development, it is conceivable that our model could directly generalize these pattern formation-based approaches.

Perhaps more closely related to our model is the ‘relaxed beliefs under psychedelics’ (REBUS) model, which proposes to explain the effects of classical psychedelics in terms of predictive coding theory [69]. Similar to the Wake-Sleep algorithm, predictive coding theory [105] models sensory representation learning with neural dynamics and local synaptic modifications that collectively optimize an ELBO objective function. However, at a mechanistic level, there are numerous differences, the most easily distinguishable feature being that the Wake-Sleep algorithm *requires* periods of offline ‘generative replay’ to train bottom-up synapses in its network, whereas predictive coding learning occurs concomitantly with stimulus presentation. Furthermore, the REBUS model of psychedelic effects is described at a computational level, in terms of a decrease in the ‘precision-weighting of top-down priors.’ While it is more difficult to map the REBUS model directly onto cortical microcircuitry, and the hallucinatory effects of such a model have, to our knowledge, not been directly analyzed, it has been shown that the proposed mechanism causes an *increase* in bottom-up information flow between cortical areas [106], in direct contrast to the effects that we have shown in our model (Figure 5c-d); there is some evidence supporting this idea [107], but noninvasive imaging studies are inconsistent on this question, with many studies showing by contrast an increase in top-down functional connectivity caused by classical psychedelic administration [99, 100], and with invasive recordings showing a decrease in the influence of bottom-up inputs [85, 86, 87]. Because interareal causal influence can be difficult to analyze statistically due to dense recurrent connectivity (i.e. correlation does not imply causation), we stress that it would be more effective to distinguish between the REBUS model and our ‘oneirogen hypothesis’ by performing direct interventions on inputs to the apical and basal dendritic compartments of pyramidal neurons in cortex, and by exploring whether psychedelic drugs affect the same circuitry that induces ‘generative replay’ during periods of sleep and quiescence. More consistent with our model, a recent non-mechanistic approach based on the DeepDream algorithm has been used to generate realistic hallucinations via increased influence from a top-down learning signal [95]; however, this model proposes no relationship between psychedelics and replay during sleep.

Lastly, it should be noted that the Wake-Sleep algorithm and our choice of network architecture constitute one particular model within a family of related models, all of which satisfy our key criteria for a good model of the ‘oneirogen hypothesis,’ namely that 1) the model has well-defined top-down and bottom-up pathways, 2) it learns a generative model of incoming sensory inputs, and 3) it uses periods of offline replay for learning through local synaptic plasticity. For example, in the Supplemental Materials, we have replicated all of our essential results for two alternative network architectures, also learned via the Wake-Sleep algorithm: one model uses within-layer recurrence to improve generative performance, while the other model uses a simpler single compartment neuron model. Furthermore, the closely related Contrastive Divergence learning algorithm for Boltzmann Machines [108] also involves alternations between Wake and generative Sleep phases, learns through local synaptic plasticity, and has been used to model hallucination disorders like Charles Bonnet Syndrome [109], though Boltzmann machines are computationally more cumbersome to train and require more non-biological network features than the Wake-Sleep algorithm. We feel as though it is important to recognize that models that satisfy these three criteria are more similar than they are different, and that it may be quite difficult to experimentally distinguish between them.

### Limitations

While our model is capable of capturing several effects of classical psychedelics, it also has several clear limitations. Firstly, while we have been able to model complex hallucination phenomena with backpropagation-trained networks, hallucinations generated by Wake-Sleep-trained networks were generally simpler, likely because the Wake-Sleep algorithm is well-known to be a less effective representation learning and generative modeling algorithm than backpropagation [35], despite its superior biological realism. This suggests that while it is quite possible for generative modeling approaches to produce complex hallucinations through non-biological means, algorithmic or architectural improvements may be necessary in order to make the performance of the more plausible Wake-Sleep algorithm closer to that achieved by state-of-the-art models.

Our model also oversimplifies several aspects of biology. In particular, we do not use neurons that respect Dale’s law [110, 111], and the majority of our efforts to map the Wake-Sleep algorithm onto biology focus on excitatory pyramidal neurons. Furthermore, though we do observe that neural dynamics can tolerate a significant amount of top-down input before disrupting perception, experiments and theoretical studies have shown that inputs to apical dendrites of pyramidal neurons do play an important role in waking perception [43, 98, 112], and are not *just* learning signals. We focused on clear distinctions between basally-driven Wake modes and apically-driven Sleep modes during training for computational efficiency reasons, and also due to the fact that parameter sharing across inference and generative networks in the Wake-Sleep algorithm is theoretically under-explored (though it is supported in closely related predictive coding approaches [105] and Boltzmann machines [108]). Future elaborations on our model could incorporate feedback control [113], attention [114], or multimodal sensory inputs [115] into top-down projections; such inputs could help explore how psychedelic hallucinations interact with attentional or feedback control systems in the brain, and have been shown to interact constructively with top-down learning signals in prior models [116, 117, 118]. Our use of VDVAEs is a positive step in this direction, but ideally such network architectures would be made compatible with the Wake-Sleep algorithm.

Lastly, our modeling focus has been exclusively on cortical plasticity and hallucination effects: it should be noted that our model has little bearing on other important features of the psychedelic experience of potential therapeutic relevance, because we have not included the effects of psychedelics on subcortical structures including the serotonergic system [119], which plays an important role in regulating mood and may be where psychedelics exert some of their antidepressant effects. Many studies of the effects of psychedelics on fear extinction focus on the hippocampus or the amygdala [120, 121, 122, 123]. These areas receive extensive innervation directly from serotonergic synapses originating from the dorsal raphe nucleus, which have been shown to play an important role in emotional learning [124]; because classical psychedelics may play a more direct role in modulating this serotonergic innervation, it is possible that fear conditioning results (in addition to the anxiolytic effects of psychedelics) cannot be attributed to a shift in balance between apical and basal synapses induced by psychedelic administration.

## Conclusions

Here we have proposed a hypothesis for the mechanism of action of psychedelic drugs in terms of its excitatory effects on the apical dendrites of pyramidal neurons, which we propose pushes network dynamics into a state normally reserved for offline replay and learning; we have also proposed a number of testable predictions which could be used to validate or invalidate our hypothesis. If validated, our model would describe a mechanism by which psychedelic drug administration causes ordinary sensory perception to become literally more dream-like; it further suggests that the plasticity increases observed during both sleep and psychedelic experience could occur via a common mechanism dedicated to sensory representation learning in the brain. Beyond classical psychedelics, further studying the balance between apical and basal dendritic inputs to pyramidal neurons in connection to replay during sleep may be relevant for explaining the hallucinatory effects of other drugs (such as ketamine) or mental disorders like schizophrenia [125].

## 4 Methods

### Model architecture and training

To model the effects of psychedelics on neural network dynamics and plasticity, we first constructed a simple model of the early visual system by training neural networks on two different image datasets (MNIST [62] and CIFAR10 [63]). Networks were trained with the Wake-Sleep algorithm [33], which requires, for each layer, two modes of stochastic network activity: a ‘generative mode’, and an ‘inference mode’. For the ‘inference’ mode we must specify a probability distribution *b*(**r**^(*l*)^ | **r**^(*l−*1)^), while for the ‘generative’ mode we must specify a separate distribution *p*(**r**^(*l*)^ | **r**^(*l*+1)^)^2^. Notice here that activity in ‘inference’ mode is conditioned on ‘bottom-up’ network states (**r**^(*l−*1)^), while activity in generative mode is conditioned on ‘top-down’ network states (**r**^(*l*+1)^) (Figure 1a).

The ‘inference mode’ specifies a probability distribution over neural activity, conditioned on the next-lower layer (where the lowest layer is the stimulus layer, i.e. **r**^(0)^ = **s**)—mechanistically it corresponds to activity generated by feedforward projections. To increase the expressive power of our neural units, we use multicompartmental neuron models similar to [126] with *N*_*d*_ dendritic compartments, whose voltages are summed nonlinearly to form the full input to the basal dendrites. For *l >* 0, layer activity is sampled from the distribution **r**^(*l*)^ *∼𝒩* (*h*(**r**^(*l−*1)^),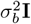), where for neuron *i* in layer *l, h*_*i*_(**r**^(*l−*1)^) is given by:

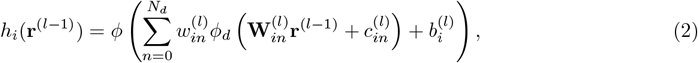

where 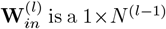 matrix of synaptic weights onto dendrite *n, c*_*in*_ is the corresponding bias for the *n*th dendritic compartment, 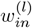 is the strictly positive weight given to the *n*th dendritic branch (roughly corresponding to a conductance), and 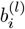 is the bias for the entire basal compartment. *ϕ*_*d*_(·) and *ϕ*(·) are nonlinearities for the dendritic branches and the total basal compartment, respectively: both are the sequential composition of the tanh nonlinearity, followed by batch normalization [127]. For the dendritic branch nonlinearities, we allow for learnable affine parameters (scale and bias), but for the entire basal dendritic compartment we constrain activity to be zero-mean and unit variance across batches in order to prevent indeterminacy between apical and basal scale parameters. For the final inference layer **r**^(*L*)^, as in the variational autoencoder [34], we parameterize both the mean *and* a diagonal covariance matrix of the inference distribution: **r**^(*L*)^ *∼𝒩 h*(**r**^(*L−*1)^), diag(*h*_2_(**r**^(*L−*1)^)), where *h*_2_(·) is also a multicompartmental model, in this case replacing the final batch normalization with an exponential nonlinearity to ensure positivity.

The ‘generative’ mode specifies a probability distribution over neural activity, conditioned on the next-higher layer—it corresponds mechanistically to activity generated by feedback projections. The highest layer, **r**^(*L*)^ is sampled from an *N* ^(*L*)^-dimensional independent standard normal distribution, **r**^(*L*)^ *∼𝒩* (0, **I**), and all subsequent layers are sampled from the distribution **r**^(*l*)^ *∼𝒩* (*µ*(**r**^(*l*+1)^),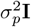), where for the *i*th neuron, *µ*_*i*_(**r**^(*l*+1)^) is given by:

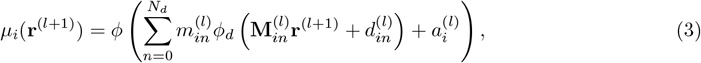

where 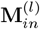 is a 1 *× N* ^(*l*+1)^ matrix of synaptic weights onto apical dendritic branch *n*, 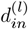 is the corresponding bias for the *n*th dendritic compartment, 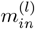 is the strictly positive weight given to the *n*th dendritic branch, and 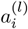 is the bias for the entire apical compartment. Again, *ϕ*_*d*_(·) and *ϕ*(·) are nonlinearities, identical to the inference (basal) pathway.

While the neuron model used here is more complicated than is normally used for single-unit neuron models, functions of this kind could feasibly be implemented by nonlinear dendritic computations [126]; we further found that using this nonlinearity qualitatively improved generative performance (Supplemental Figure S2). Given these parameterized probability distributions, we then determined the neural activity for each layer *l* according to Eq. 1. Our network trained on MNIST was composed of 3 layers, with widths [32, 16, 6], listed in ascending order. A full list of network hyperparameters for both our MNIST and CIFAR10-trained networks can be found in the Supplemental Methods.

All synaptic weights and parameters in our networks were trained via the Wake-Sleep algorithm [33], which is known to produce ‘local’ parameter updates for a wide range of neuron models (and rate or spike-based output distributions), though the specific functional form of the update may vary depending on the neuron model chosen [128]. These updates, for reasonable choices of neural network architecture, can be interpreted as predictions for how synaptic plasticity should look in the brain, if learning were really occurring via the Wake-Sleep algorithm or some approximation thereof.

Consider a generic inference (basal dendrite) parameter for neuron *i*,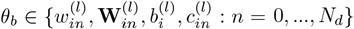. The Wake-Sleep algorithm gives the following update, for a single stimulus presentation:

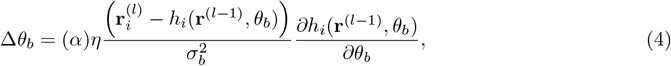

where *η* is a learning rate, and the gate *α* ensures that learning only occurs during sleep mode. Further, for reasons of computational efficiency, we average weight updates across a batch of 512 stimulus presentations; similar results could in principal be obtained with purely online updates [129], but we opted to present stimuli in batches here in order to parallelize computations. 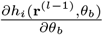 changes depending on the parameter *θ*, reflecting that particular parameter’s contribution to basal dendritic activity. For a dendritic branch weight 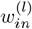, we have:

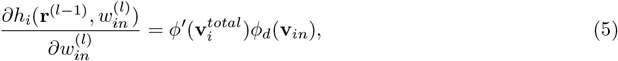

where 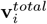 is the total input to the basal dendritic compartment, and 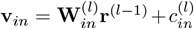 is the total input to the *n*th dendritic branch. This update has the functional form of a classical ‘delta’ learning rule [88], where a compartmental prediction error between local dendritic activity and neuronal firing rate is multiplicatively combined with branch-specific input to provide changes in the conductance for the *n*th branch. Similarly, for the *j*th synapse on the *n*th dendritic branch, 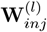, we have:

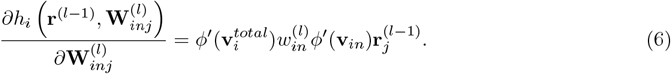

Unlike for simple one-compartment neuron models, computation of parameter updates for dendritic synapses 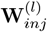 requires weighting the ‘delta’ error by the conductance of the corresponding dendritic branch (*w*_*in*_), which could be approximated by the passive diffusion of signaling molecules from the principal basal dendritic compartment back along dendritic branches to individual synapses.

For generative parameters 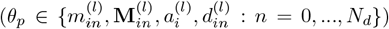, we have a nearly identical update for a single stimulus presentation:

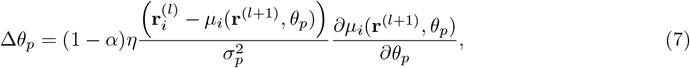

where now input in the apical dendritic compartment, *µ*_*i*_(**r**^(*l*+1)^), is being compared to the activity of the neuron as a whole to determine the magnitude and sign of plasticity. The (1 *− α*) gate in this case ensures that plasticity only occurs during the Wake mode. We provide pseudocode (Algorithm 1) for our Wake-Sleep implementation, as well as a full list of algorithm and optimizer hyperparameters (Tables S1 and S2) in the Supplemental Materials^3^.

### Modeling Hallucinations

During training, neural network activity is either dominated entirely by bottom-up inputs (Wake, *α* = 0) or by top-down inputs (Sleep, *α* = 1). As a consequence, sampling neural activity is computationally low-cost, and can be performed in a single time step. During Wake, one can take a sampled stimulus variable **s**, determine the activity at layer 1, then 2, and so on until layer *L*, while during Sleep, one can sample a latent network state in layer *L* and traverse the layers in reverse order, down to the stimulus layer. However, this is not possible if *α ∉ {*0, 1*}*, because activity in each layer *l* should depend simultaneously on layer *l* + 1 and layer *l−*1. For this reason, we chose to model hallucinatory neural activity *dynamically*, as follows:

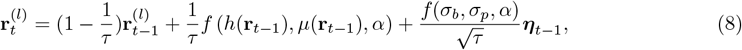

where *τ* is a time constant that determines how much of the previous network state is retained, and ***η****t−*1 *∼𝒩* (0, **I**). Critically, if we take *τ* = 1 these dynamics reduce to the sampling procedure used during training (Eq. 1). *A priori*, the choice of interpolation function *f* (*a, b, α*) is arbitrary. We selected the following function:

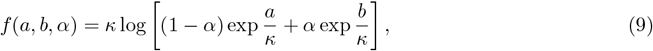

where *κ* = 0.35 is a free parameter. This function is equivalent to linear interpolation as *κ → ∞*, and is equivalent to the maximum function between arguments *a* and *b* as *κ →* 0 if *α* = 0.5. By selecting *κ* = 0.35, we are biasing the system towards registering positive inputs from apical *or* basal sources (in the inclusive sense). We found that this produced ‘hallucinatory’ percepts in stimulus space that did not reduce the intensity of input stimuli as *α* increased; rather, inputs maintained their intensity and hallucinations were added on top if they were of greater intensity than the ground-truth image. All simulations were run for 800 timesteps, with *τ* = 0.1. As a control, we compared our results to network dynamics produced purely by increases in noise, without increases in apical dendritic influence (which we refer to as our noise-based hallucination protocol). For these control simulations, we produced network activity time series with the following equation:

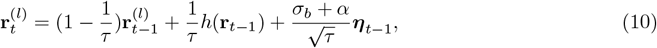

so that the standard deviation of the injected noise increased linearly with *α*.

### Apical and Basal Alignment

To measure the alignment between inputs in the apical and basal dendritic compartments of our model neurons, we computed the ‘Wake’ neural responses to the full test dataset and measured the activity in both the basal and apical compartments of our neurons (*h*(**r**^(*l−*1)^) and *µ*(**r**^(*l*+1)^), respectively). We then calculated the correlation coefficient between apical and basal compartments for the same neuron, compared to the correlation between compartments for two randomly selected neurons.

### Quantifying plasticity

To quantify the total amount of plasticity induced in our model system by the administration of psychedelic drugs, we measured the change in *relative* parameter strength (averaging across all synapses in the network and an ensemble of 512 test images). For each test image, we simulated network dynamics according to Eq. 8. Subsequently, for each parameter *θ*, we calculated the net amount of plasticity induced by viewing all test images, Δ*θ*. We subsequently reported the relative change:

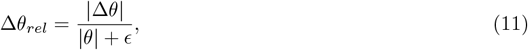

under conditions in which *α* values gate plasticity (as in ordinary Wake-Sleep) and under conditions in which psychedelic drug administration does not also affect plasticity gating. Here, we took *ϵ* = 10^*−*2^ to avoid numerical instabilities.

### Classifier training

As we trained our neural network using the Wake-Sleep algorithm, we simultaneously trained a *separate* classifier network based on Wake-phase neural activity in the second network layer on a cross-entropy loss, to identify the stimulus class of the input to the system. For our classifier, we used a multilayer perceptron neural network with a single 256-unit hidden layer and tanh(·) nonlinearities.

We then quantified the accuracy of the classifier on the test set, based on neural activity drawn from the final time step *T* of hallucination simulations with various values of *α*. We further measured the average variance of the 10-dimensional output logits of the neural network.

### Quantifying correlation matrix similarity before and after psychedelics

To quantify how similar the pairwise correlations between neurons in our model networks were before and after the administration of psychedelics, we recorded hallucinatory network dynamics for an ensemble of 512 test images, and measured pairwise correlations between neurons in the first network layer. To compare these matrices, we then report the correlation coefficient between the flattened *N × N* matrices. For this metric, a value of 1 indicates that the correlation matrices are perfectly aligned, while a value of -1 indicates that pairwise correlations are fully inverted.

### Quantifying interareal causality through inactivations

To quantify changes in interareal functional connectivity induced by psychedelics, we performed two different types of inactivation. In the first, we inactivated the apical dendritic compartments of all neurons in the stimulus layer, and measured how this inactivation affected across-stimulus variability of neurons relative to the fully active state. In the second method, we inactivated all neurons in the deepest layer, and measured the same effect in across-stimulus variability in the stimulus layer. For both inactivation schemes, we report the mean and standard error of the variance ratio:

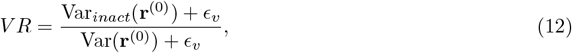

where we added *ϵ*_*v*_ = 10^*−*3^ to the denominator to prevent numerical instability and to the numerator ensure that the ratio evaluates to 1 if the two variances are equivalent.

### Generating hallucinations in hierarchical variational autoencoders

To model more complex hallucination phenomena than could be observed in our simpler Wake-Sleep-trained networks, we used pretrained VDVAE [79] models trained on Tiny ImageNet [80], a 64x64 pixel variant of ImageNet, and FFHQ-256 [81], a dataset of 256x256 pixel human faces. VDVAE models are very similar to our Wake-Sleep-trained models: they are trained on the same unsupervised representation learning objective function (the ELBO), and every layer of the multilayer network models are parameterized by a bottom-up inference distribution *b* and a top-down generative distribution *p*. VDVAE models are top-down VAEs [77], which means that the inference distribution is conditioned on bottom-up stimuli *and* latent network activity at higher layers, i.e. the distribution is written *b*(**r**^(*l*)^ | *h*(**s, r**^(*l*+1)^)), where *h*(·) is a parameterized neural network. By contrast the generative distribution is conditioned only on top-down inputs, and is written *p*(**r**^(*l*)^ | *µ*(**r**^(*l*+1)^)), where *µ*(**r**^(*l*+1)^) is also a parameterized neural network.

For our Wake-Sleep-trained networks, we modeled hallucinations by simulating a stochastic time series at each layer (Eq. 8), but for the VDVAE models we found this to be computationally infeasible. Instead, we modeled hallucinations with a single bottom-up and top-down pass through the network, as follows:

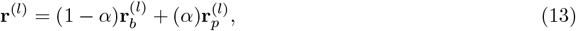

where 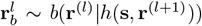 is a sample from the inference distribution, and 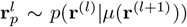 is a sample from the generative distribution. This generation scheme is simpler and less computationally expensive than our previous method, while still producing purely Wake-stage sampling when *α* = 0 and Sleep-stage sampling when *α* = 1; intermediate values of *α* correspond to modeled hallucinatory network states^4^. Our Laplacian pyramid analysis of generated images was performed using the Pyrtools package [130].

## Supporting information

Figure 2a multicompartmental mnist hallucination video

Figure 2c multicompartmental cifar10 hallucination video

## Acknowledgements and Funding

We would like to thank members of both G.L. and B.R.’s labs, as well as James M. Shine, Brandon Munn, Christopher Whyte, Veronica Chelu, Jiameng Wu, Matthew Larkum, Santiago Jaramillo, Michael Wehr, Neil Savalia, Alexandra Klein, Sarah Cook, Conor Lane, Anousheh Bakhti-Suroosh, Runchong Wang, Michael Okun, and Jordan O’Byrne for insightful discussions and feedback. This work was supported by: [GL] NSERC Discovery Grant (RGPIN-2018-04821), Canada CIFAR AI Chair Program, Canada Research Chair in Neural Computations and Interfacing (CIHR, tier 2). [BR] NSERC (Discovery Grant: RGPIN-2020-05105; Discovery Accelerator Supplement: RGPAS-2020-00031; Arthur B. McDonald Fellowship: 566355-2022) and CIFAR (Canada AI Chair; Learning in Machine and Brains Fellowship). [CB] is supported in part by the FRQNT Strategic Clusters Program (Centre UNIQUE - Quebec Neuro-AI Research Center). The authors acknowledge the material support of NVIDIA in the form of computational resources.

## 6 Ethics Declarations

Psychedelic drug research has a long history fraught with many instances of unethical research practice [131]. Further, psychedelic drug use itself has long been stigmatized and punished through legal measures [132], often at the expense of indigenous peoples, who have long incorporated psychoactive substances into their cultural and spiritual practices [133]. In the interest of avoiding a repetition of past mistakes, we feel compelled to provide explicit guidance on how our work should be interpreted and used. To do so, we will take inspiration from two principal ethical frameworks: the Montreal Declaration on Responsible AI [134], and the EQUIP framework for equity-oriented healthcare [135, 136]. We strongly encourage anyone considering extending our research or using our work in any form of clinical setting to ensure that subsequent research adheres to these frameworks.

Below, drawing from these ethical frameworks, we will provide a set of guidelines for how our work should be interpreted and used. Though these guidelines are by no means exhaustive, our hope is that adherence to them will help promote the potential positive outcomes of our work while limiting potential negative consequences.

### Guidelines for the ethical use of this study

#### Do

1. Ensure that the elements of our hypothesis have been adequately tested, as outlined in our discussion, *before* using our framework in any form of clinical or therapeutic setting.

2. Use our ideas to inform further basic neuroscience research on perception, learning, sleep, and replay phenomena.

3. Explore our ideas as an opportunity to inform your own understanding of cognition, learning, and perception, with the understanding that these ideas have not yet been fully validated experimentally.

4. Feel free to ask us if you are worried that your proposed use of our work may have negative impacts.

#### Do not

1. Report our results as scientific fact. We have outlined a *hypothesis*, which is designed to be tested by the experimental neuroscience community.

2. Cite or interpret our results without an adequate understanding of the evidence supporting the various claims made in this study. Feel free to ask us if you are worried that you may be misinterpreting our results.

3. Use our results to extract undue or inequitable profit. The ideas developed in this paper are the product of decades of research and public funding, built upon *centuries* of exploration of psychedelics. Any knowledge or value contained within this paper is the common heritage of all humanity, with particular recognition due to the indigenous and marginalized communities that have historically suffered and are currently suffering from oppressive government and industry policies.

4. Use our results for any application that could violate human rights or harm human beings in any way.

## A Supplementary Materials

### Recurrent network model

To explore the extent to which our results hold for different neuron models, and to give our generative model more expressive power than the traditional Helmholtz machine [137], we constructed a network model with a single timestep of within-layer recurrent denoising in each layer, which gives our model some similarities to denoising diffusion approaches [138]. For both our ‘inference’ mode and our ‘generative’ mode we specify both a denoised network state 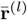 and a noise-corrupted network state **r**^(*l*)^ for layer *l*; specifying a neural network model is then equivalent to specifying, for each layer, a joint probability distribution over denoised and noise-corrupted network states for both the inference and generative modes, i.e. for the ‘inference’ mode we must specify a probability distribution 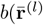, **r**^(*l*)^ |**r**^(*l−*1)^), while for the ‘generative’ mode we must specify a separate distribution 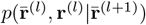 ^5^. Notice here that activity in ‘inference’ mode is conditioned on ‘bottomup’ network states (**r**^(*l−*1)^), while activity in generative mode is conditioned on ‘top-down’ network states 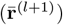 (Figure 1a).

The ‘inference mode’ specifies a probability distribution over neural activity, conditioned on the next-lower layer (where the lowest layer is the stimulus layer, i.e. **r**^(0)^ = **s**)—mechanistically it corresponds to activity generated by feedforward projections. For *l >* 0, layer activity is sampled from the distribution 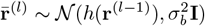), where *h*(**r**^(*l−*1)^) is given by:

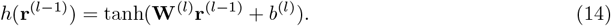

Subsequently, we add additional noise to get a noise-corrupted network state 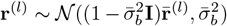; while noise corruption is a natural feature of network dynamics in the brain [139], we include it here in our model because it has been shown that *denoising* is a critical aspect of many powerful generative modeling approaches [140, 141, 142], and we have likewise found that it improves the quality of generated images in our learned networks (Supplemental Figure S2).

The ‘generative’ mode specifies a probability distribution over neural activity, conditioned on the next-higher layer—it corresponds mechanistically to activity generated by feedback projections. The highest layer, **r**^(*L*)^ is sampled from an *N* ^(*L*)^-dimensional independent standard normal distribution, **r**^(*L*)^ *∼𝒩* (0, **I**), and all subsequent layers are sampled from the distribution 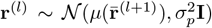, where 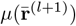 is given by:

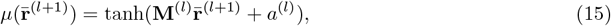

where *m*^(*l*)^ is a *N* ^*l*^ *×N* ^(*l*+1)^ weight matrix, and *a* is a bias term. Subsequently, the network goes through a single timestep of recurrent denoising, so that 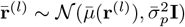, where 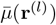 is given by:

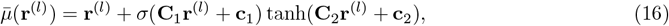

where *σ*(·) is a sigmoid nonlinearity that acts as a gating function similar those used in the LSTM [143] and GRU [144], **C**_1_ and **C**_2_ are *N* ^(*l*)^ *× N* ^(*l*)^ recurrent weight matrices, and **c**_1_ and **c**_2_ are biases. While this is a more complicated nonlinearity than is normally used for single-unit neuron models, functions of this kind could feasibly be implemented by nonlinear dendritic computations [126]; we further found that using this nonlinearity qualitatively improved generative performance. Given these parameterized probability distributions, we then determined the neural activity for each layer *l* according to Eq. (1). As with our multicompartmental neuron model, inference and generative parameters were updated according to Eqs. (4) and (7), respectively. Recurrent network hyperparameters are available in Table S3.

### Simplified neuron model

As a control, we also tested our results using a simplified multilayer perceptron neuron model, which used neither batch normalization nor multiple dendritic branches. For the ‘inference’ mode within the simplified model, for *l >* 0, layer activity is sampled from the distribution **r**^(*l*)^ *∼𝒩* (*h*(**r**^(*l−*1)^),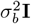), where for neuron *i* in layer *l, h*_*i*_(**r**^(*l−*1)^) is given by:

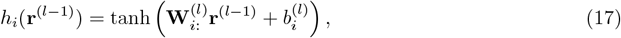

where 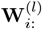 is a 1 *× N* ^(*l−*1)^ matrix of basal synaptic weights onto neuron *i*, and *b*_*i*_ is the corresponding bias.

The simplified ‘generative’ mode likewise replaces the branched neuron model used in the main text with a multilayer perceptron model. The highest layer, **r**^(*L*)^ is sampled from an *N* ^(*L*)^-dimensional independent standard normal distribution, **r**^(*L*)^ *∼𝒩* (0, **I**), and all subsequent layers are sampled from the distribution **r**^(*l*)^ *∼𝒩* (*µ*(**r**^(*l*+1)^),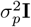), where for the *i*th neuron, *µ*_*i*_(**r**^(*l*+1)^) is given by:

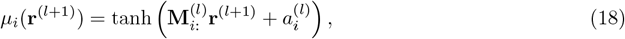

where 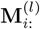 is a 1 *× N* ^(*l*+1)^ matrix of apical synaptic weights onto neuron *i*, and 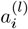 is the corresponding bias. As with the branched neuron model, inference and generative parameters were updated according to Equations (4) and (7), respectively. For optimization, we used the identical hyperparameters to the multicompartment neuron model (Table S1).

**Figure S1:**
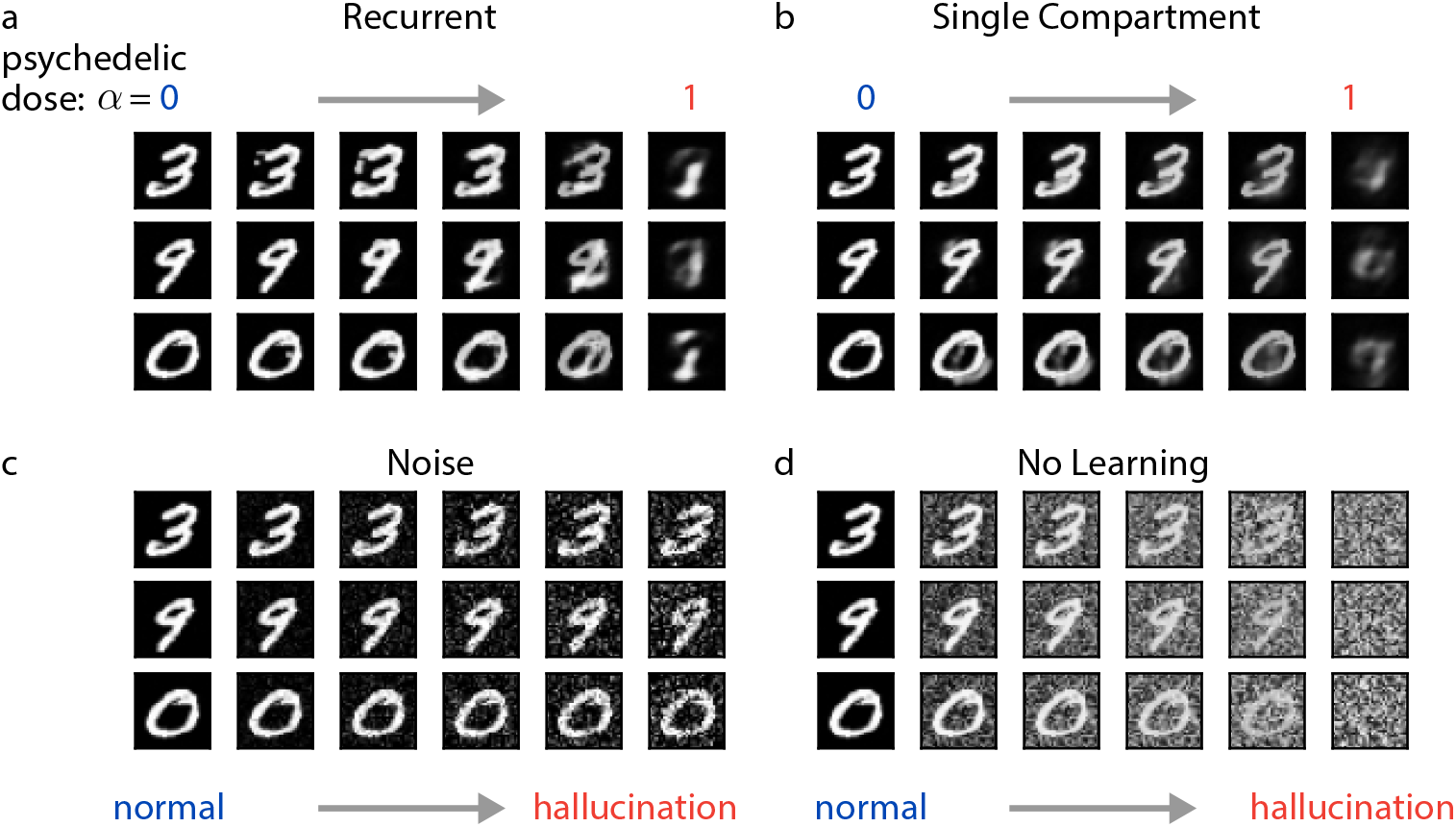
Visualizing the effects of psychedelics for alternative model architectures. We model the effects of classical psychedelics by progressively increasing *α* from 0 to 1 in alternative model architectures. We visualize the effects of psychedelics on the network representation by inspecting the stimulus layer **s. a)** Example stimulus-layer activity (rows) in response to an MNIST digit presentation as psychedelic dose increases (columns, left to right) in the recurrent network model. **b)** Same as (a) but for our single compartment neuron model. **c)** Same as (a) using the multicompartment neuron model used for our main results, but for our noise-based hallucination hallucination protocol. **d)** Same as (c), but in a network in which neither the generative nor inference pathways have been trained beyond initialization.

**Figure S2:**
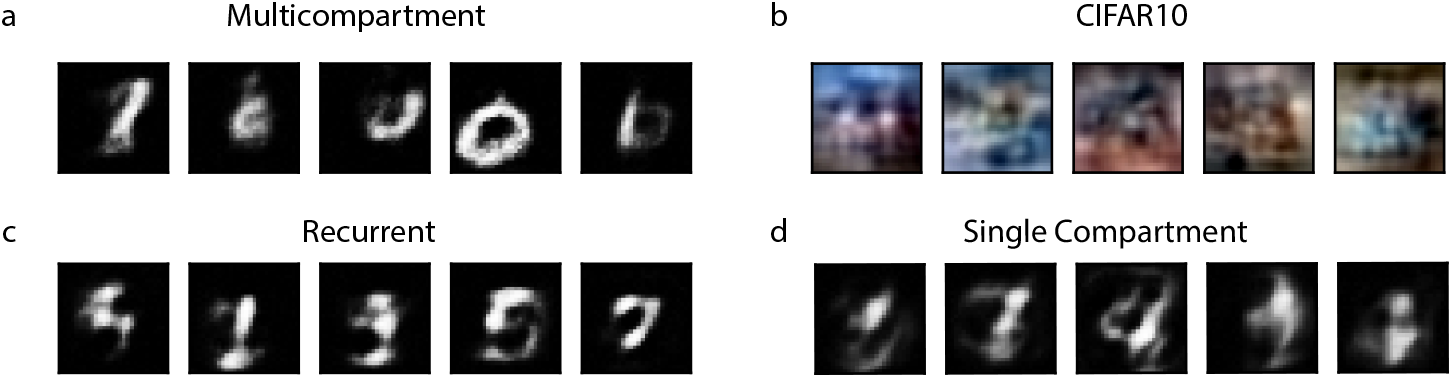
Example generated images for different model architectures and datasets. Generated images sampled from Eq. (1) with *α* = 1 for: **a)** Our primary multicompartment neuron model trained on MNIST, **b)** A multicompartment neuron model trained on CIFAR10, **c)** The recurrent network model, **d)** The single compartment neuron model.

**Figure S3:**
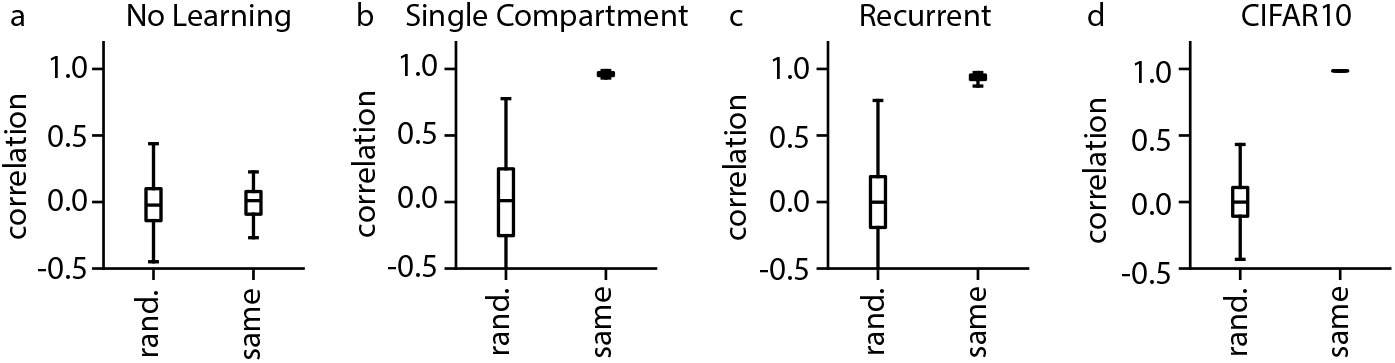
Alignment between apical and basal dendritic compartments for different model architectures and datasets. Apical-basal alignment for: **a)** An untrained multicompartment neuron model trained on MNIST, **b)** A single compartment neuron model, **c)** A recurrent network model, **d)** A multicompartment neuron model trained on CIFAR10.

**Figure S4:**
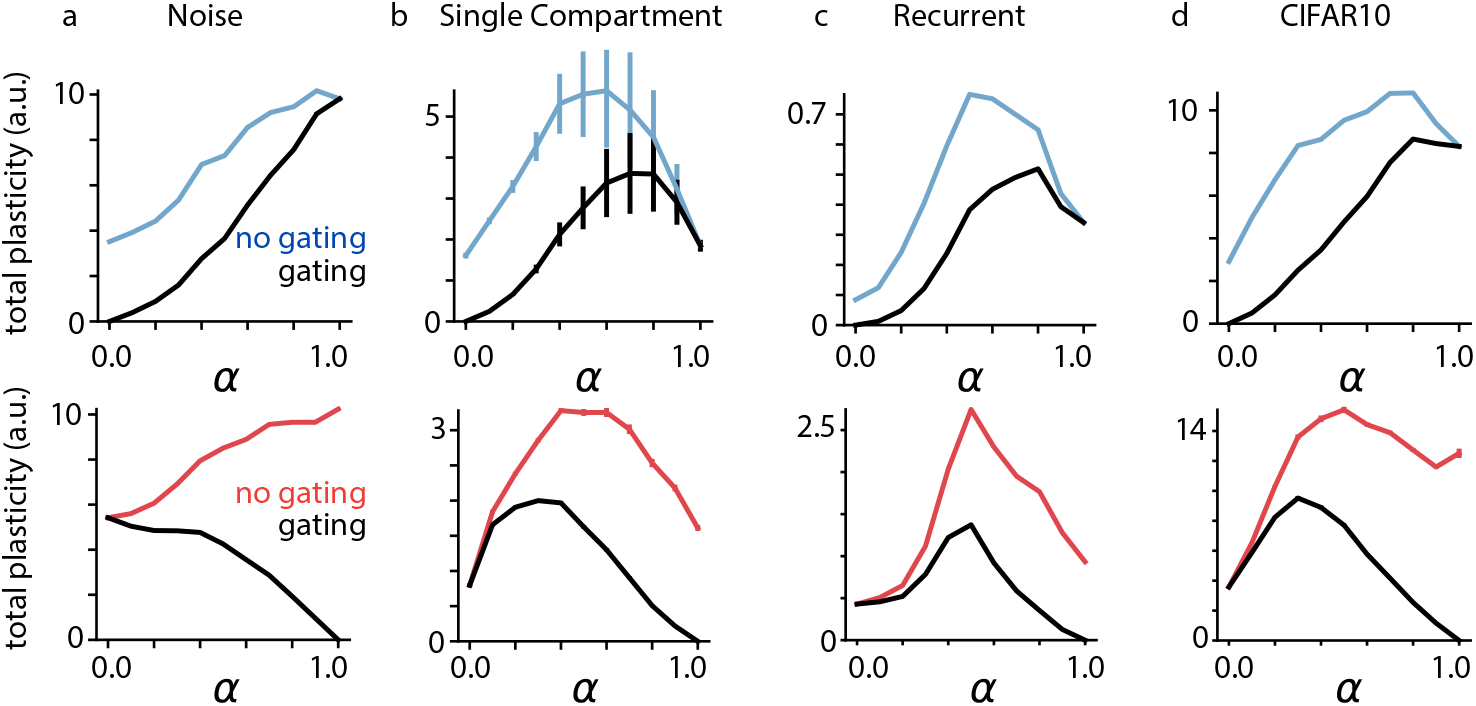
Hallucination-induced synaptic plasticity for different neuron models. **a)** Basal (top) and apical (bottom) plasticity as a function of *α* for a multicompartment neuron model trained on MNIST, using our noise-based hallucination protocol as a control. **b)** Same as (a) for a single compartment neuron model, using our primary hallucination protocol. **c)** Same as (b) for a recurrent network model, **d)** Same as (b) for a multicompartment neuron model trained on CIFAR10. Error bars indicate +/-1 s.e.m.

**Figure S5:**
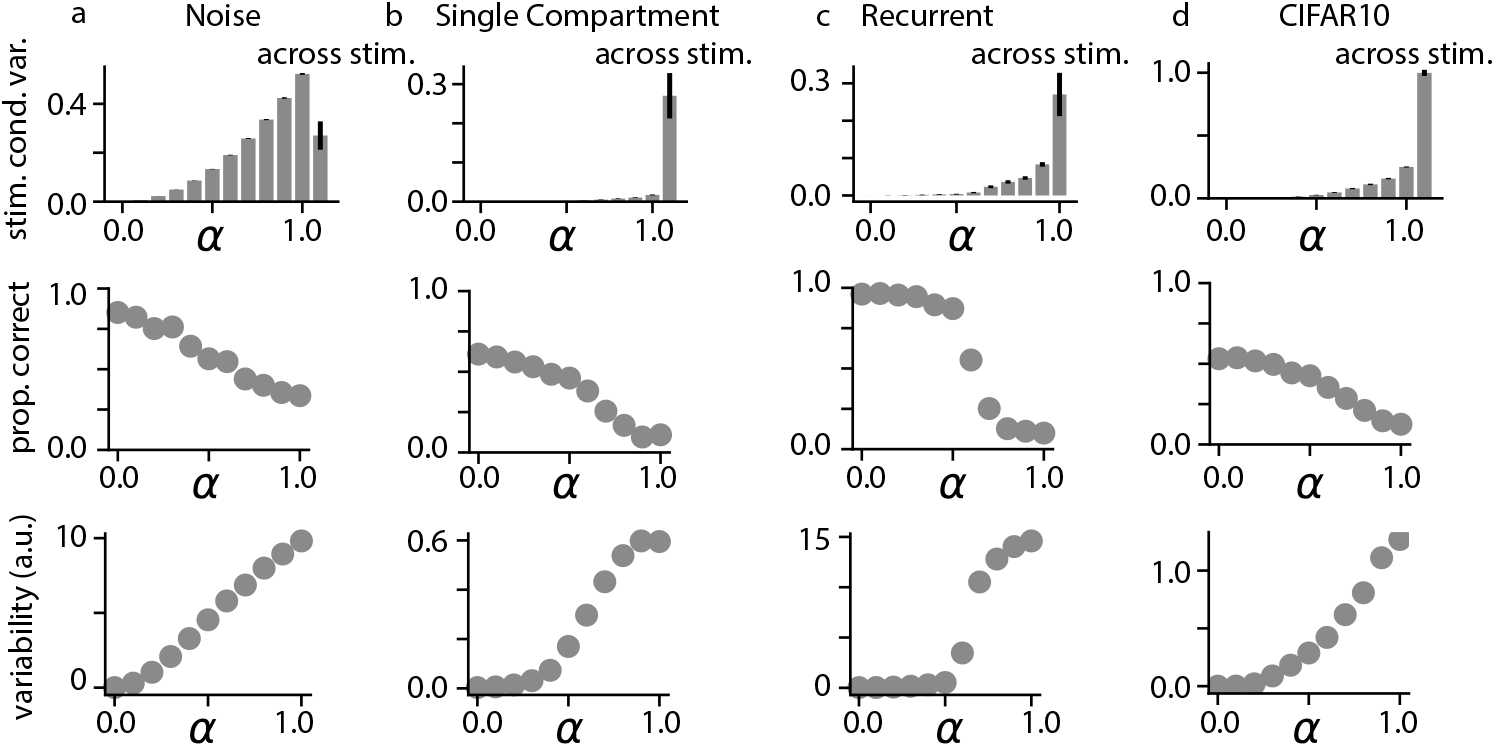
Neural variability changes for different neuron models. **a)** Stimulus-conditioned variability (top), classifier accuracy (middle), and classifier output variability (bottom) as a function of *α* for a multicompartment neuron model trained on MNIST, using our noise-based hallucination protocol as a control. **b)** Same as (b) for a single compartment neuron model, using our primary hallucination protocol. **c)** Same as (b) for a recurrent network model, **d)** Same as (b) for a multicompartment neuron model trained on CIFAR10. Error bars indicate +/-1 s.e.m.

**Figure S6:**
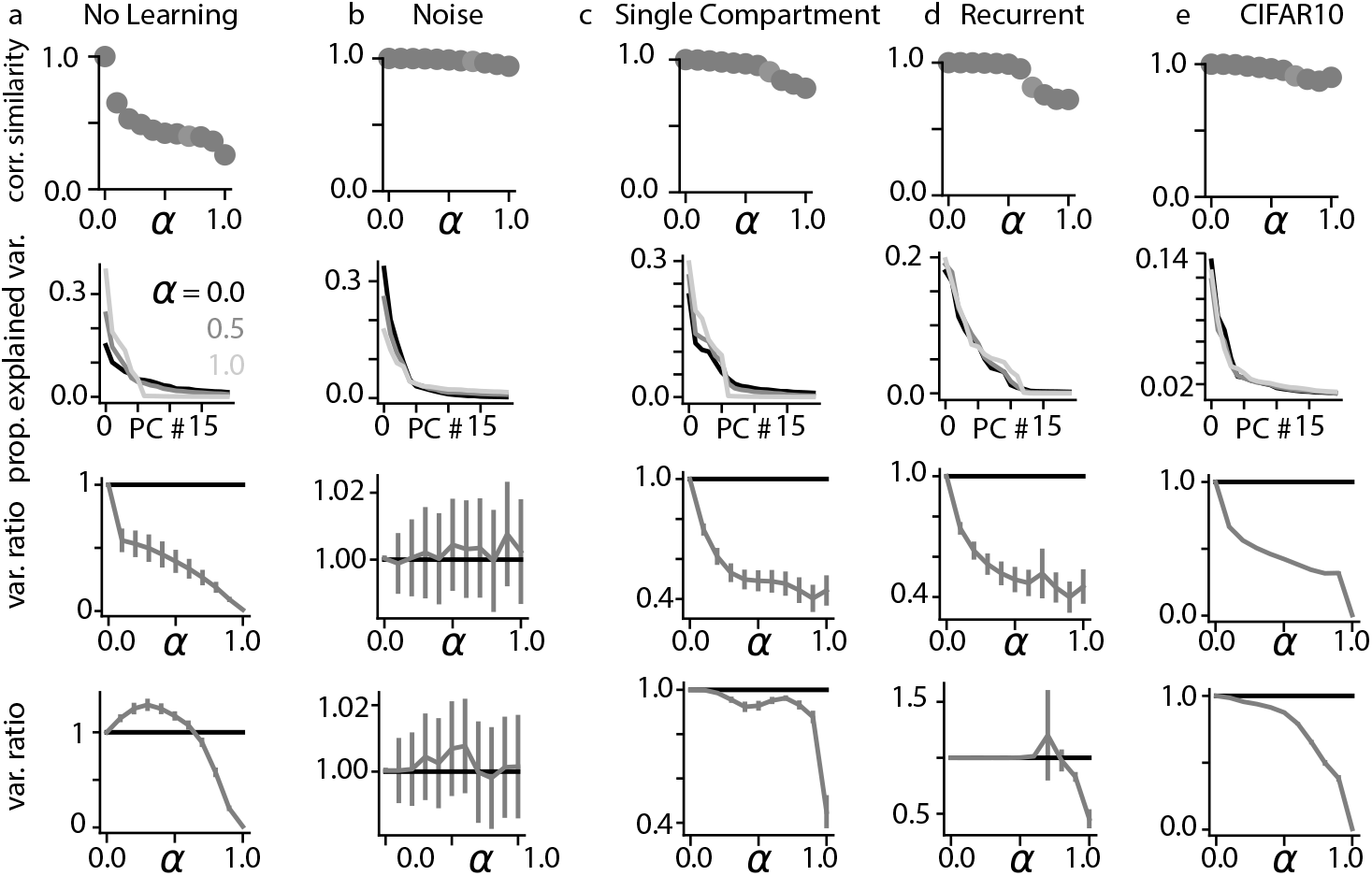
Network-level effects of psychedelics for different network architectures and training datasets. For each network architecture, we examine: correlation similarity as a function of *α* (top row), the proportion explained variance across stimuli as a function of principal component number (second row), the ratio of across-stimulus variance in stimulus layer neurons when apical dendrites have been inactivated compared to baseline conditions across different *α* values (third row), and the ratio of across-stimulus variance in stimulus layer neurons when the deepest network layer has been inactivated across different *α* values (fourth row). **a)** Results for an untrained multicompartment neuron. **b)** Results for a multicompartment neuron model trained on MNIST, using our noise-based hallucination protocol. **c)** Results for a single compartment neuron model. **d)** Results for a recurrent network model. **e)** Results for a multicompartment neuron model trained on CIFAR10. Error bars indicate +/-1 s.e.m.

**Figure S7:**
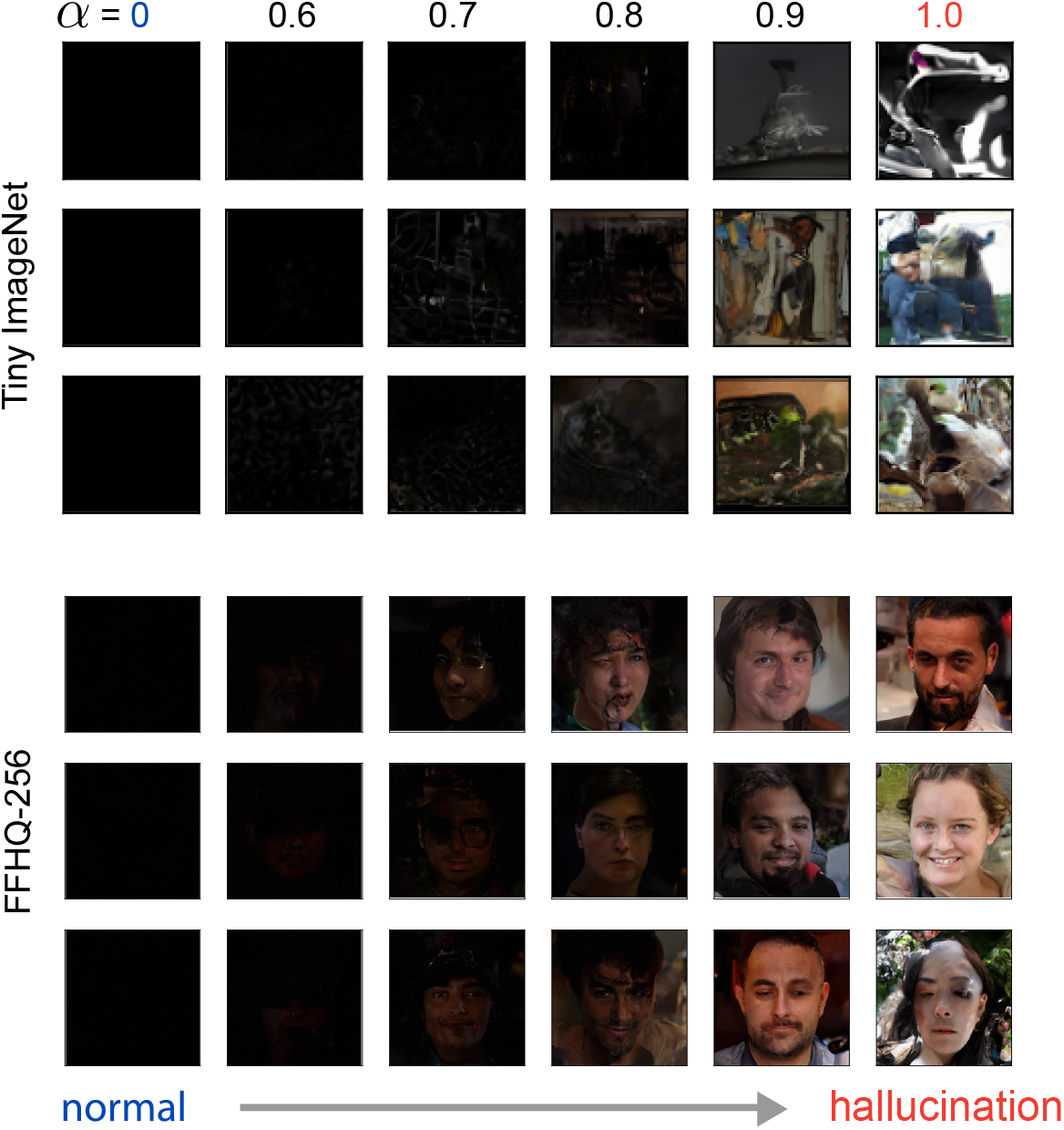
Visualizing the eyes-closed effects of psychedelics in pretrained VDVAE models. Decoded outputs of a pretrained VDVAE model trained on Tiny ImageNet (Top) and FFHQ-256 (Bottom) based on hallucinations generated in the top 35 layers of the model. Black input images were used for samples in all rows. Hallucination intensity, parameterized by *α*, increases along columns.

**Figure S8:**
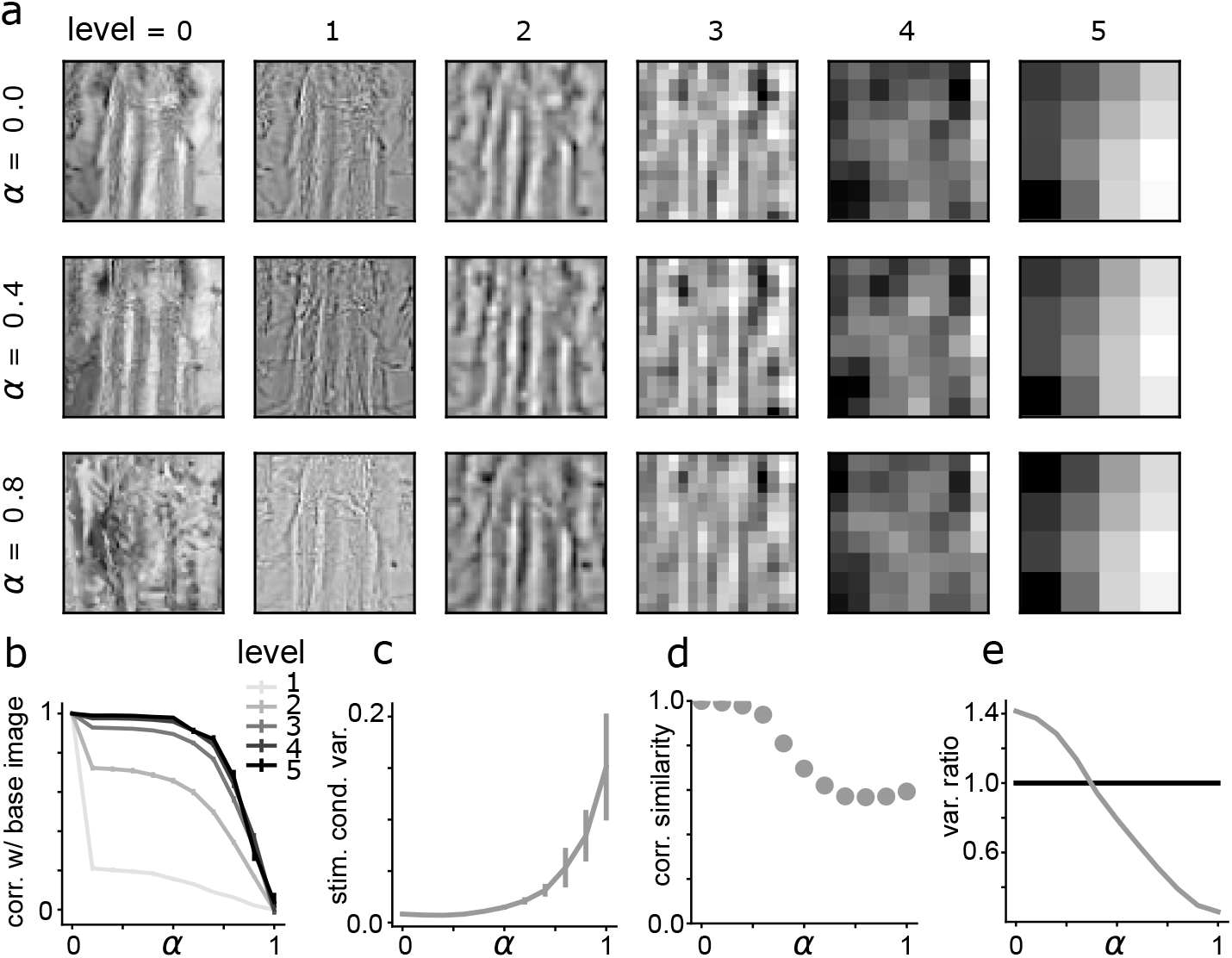
Analyzing the image- and network-level effects of psychedelics in a Tiny ImageNet-pretrained VDVAE model. **a)** Laplacian pyramid features for a grayscale example input image from the Tiny ImageNet dataset (top left). Pyramid levels increase along columns, corresponding to decreasing resolution, and hallucination intensity increases along rows. **b)** Correlation similarity across pyramid levels between the base image and a hallucinated image across different *α* values, averaged over 100 image samples. **c)** Stimulus-conditioned variance of units in layer 30 (descending from the top of the network) across different *α* values, averaged over 100 sample images and 32 distinct trials. **d)** Correlation similarity calculated between correlation matrices for units in layer 30 across different *α* values, averaged over spatial positions and 100 sample images. **e)** Ratio of across-stimulus variance in individual units of layer 30 when the highest 20 layers of the network have been inactivated, versus baseline conditions across different *α* values. Error bars indicate +/-1 s.e.m.

**Table S1:**
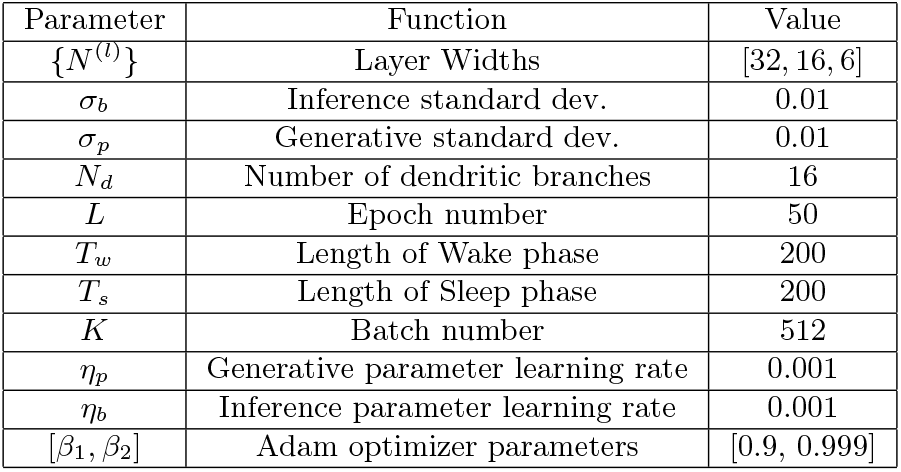
MNIST multicompartment network hyperparameters.

| Parameter | Function | Value |
| --- | --- | --- |
| $\{N^{(l)}\}$ | Layer Widths | [32, 16, 6] |
| $\sigma_b$ | Inference standard dev. | 0.01 |
| $\sigma_p$ | Generative standard dev. | 0.01 |
| $N_d$ | Number of dendritic branches | 16 |
| $L$ | Epoch number | 50 |
| $T_w$ | Length of Wake phase | 200 |
| $T_s$ | Length of Sleep phase | 200 |
| $K$ | Batch number | 512 |
| $\eta_p$ | Generative parameter learning rate | 0.001 |
| $\eta_b$ | Inference parameter learning rate | 0.001 |
| $[\beta_1, \beta_2]$ | Adam optimizer parameters | [0.9, 0.999] |

**Table S2:**
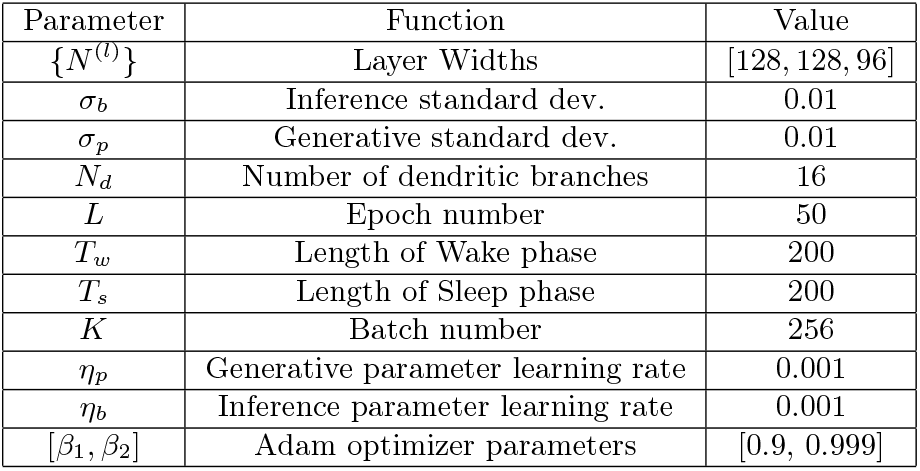
CIFAR10 multicompartment network hyperparameters.

**Table S3:** Recurrent network hyperparameters.

| Parameter | Function | Value |
| --- | --- | --- |
| $\{N^{(l)}\}$ | Layer Widths | [512, 256, 256, 128, 64, 32, 16] |
| $\sigma_b$ | Denoised inference standard dev. | 0.01 |
| $\bar{\sigma}_b$ | Noisy inference standard dev. | 0.01 |
| $\sigma_p$ | Denoised generative standard dev. | 0.01 |
| $\bar{\sigma}_p$ | Noisy generative standard dev. | 0.01 |
| $L$ | Epoch number | 400 |
| $T_w$ | Length of Wake phase | 200 |
| $T_s$ | Length of Sleep phase | 50 |
| $K$ | Batch number | 512 |
| $\eta_p$ | Generative parameter learning rate | 0.001 |
| $\eta_b$ | Inference parameter learning rate | 0.0001 |
| $[\beta_1, \beta_2]$ | Adam optimizer parameters | [0.9, 0.999] |

#### Algorithm 1 Wake-Sleep Pseudocode

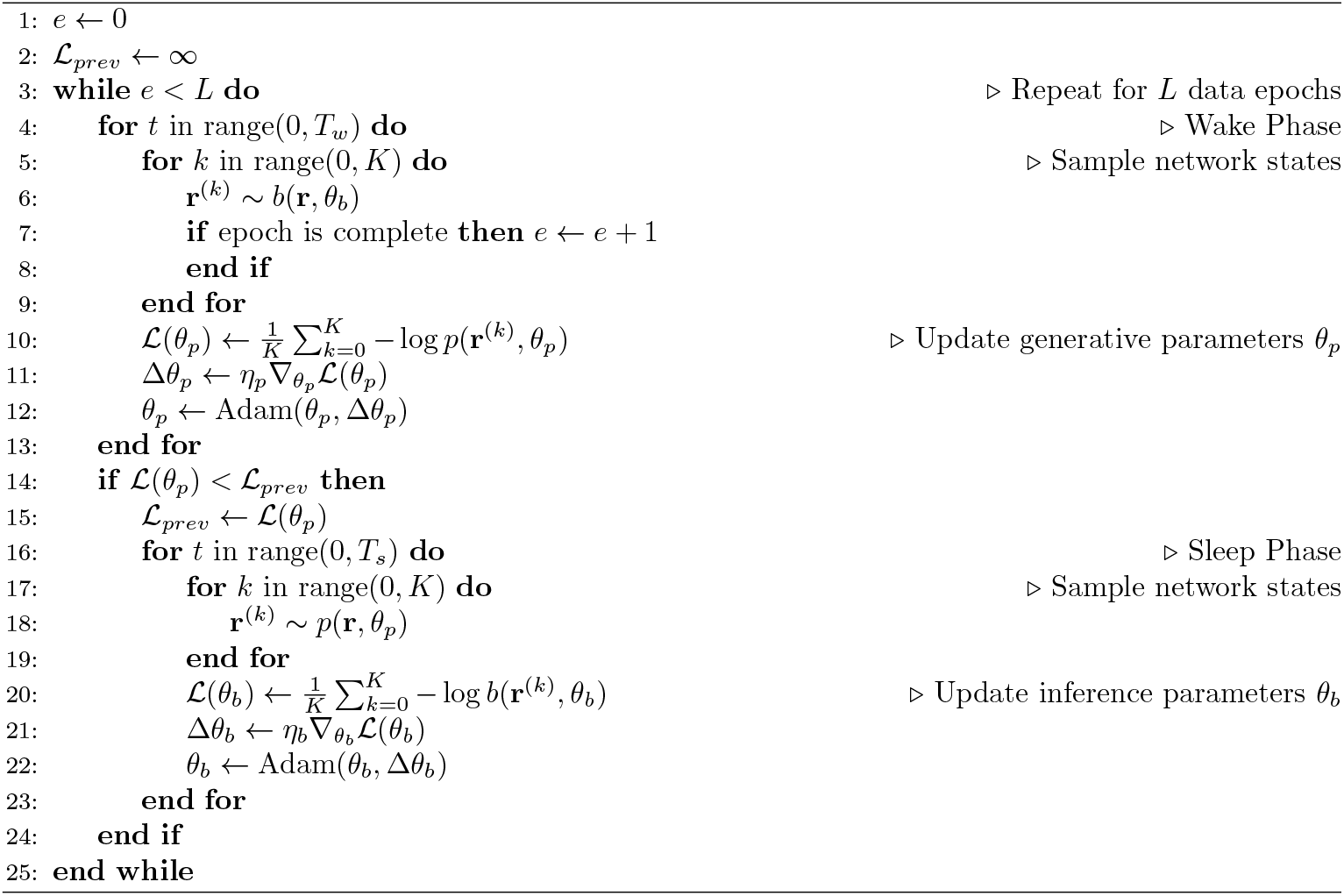

## Footnotes

1 For readers who are unfamiliar with the Wake-Sleep algorithm, a tutorial can be found here [42].

2 As a notational convention, we will use letters when referring to mathematical objects from the generative, top-down distribution, and their vertical reflection when referring to the inference, bottom-up distribution (e.g. *p* and *b*)

3 Code for reproducing all results from Wake-Sleep-trained models this study is available here: https://github.com/colinbredenberg/oneirogen-hypothesis.

4 Code for reproducing results obtained with pretrained VDVAE models is available here: https://github.com/colinbredenberg/vdvae.

5 As a notational convention, we will use letters when referring to mathematical objects from the generative, top-down distribution, and their vertical reflection when referring to the inference, bottom-up distribution (e.g. *p* and *b*)

## Notes

### Competing Interest Statement

The authors have declared no competing interest.

### Summary of Updates

Figures 6, S7, S8 added; results and methods updated with new experiments exploring modeled hallucinations in Very Deep Variational Autoencoder models; text updated to improve the quality of the literature review.

